# Changes in social behaviour with alterations of MAPK3 and KCTD13/CUL3 pathways in two new outbred rat models for the 16p11.2 syndromes with autism spectrum disorders

**DOI:** 10.1101/2023.02.11.528123

**Authors:** Sandra Martin Lorenzo, Maria Del Mar Muniz Moreno, Helin Atas, Marion Pellen, Valérie Nalesso, Wolfgang Raffelsberger, Geraldine Prevost, Loic Lindner, Marie-Christine Birling, Séverine Menoret, Laurent Tesson, Luc Negroni, Jean-Paul Concordet, Ignacio Anegon, Yann Herault

**Author notes:** **Correspondence:** Yann Herault.

## Abstract

Copy number variations (CNVs) of the human 16p11.2 locus are associated with several developmental/neurocognitive syndromes. Particularly, deletion and duplication of this genetic interval are found in patients with autism spectrum disorders, intellectual disability and other psychiatric traits. The high gene density associated with the region and the strong phenotypic variability of incomplete penetrance, make the study of the 16p11.2 syndromes extremely complex. To systematically study the effect of 16p11.2 CNVs and identify candidate genes and molecular mechanisms involved in the pathophysiology, mouse models were generated previously and showed learning and memory, and to some extent social deficits. To go further in understanding the social deficits caused by 16p11.2 syndromes, we engineered deletion and duplication of the homologous region to the human 16p11.2 genetic interval in two rat outbred strains, Sprague Dawley (SD) and Long Evans (LE). The 16p11.2 rat models displayed convergent defects in social behaviour and only a few cognitive defects. Interestingly major pathways affecting MAPK3 and CUL3 were found altered in the rat 16p11.2 models with additional changes in males compared to females. Altogether, the consequences of the 16p11.2 genetic region dosage on social behaviour are now found in three different species: humans, mice and rats. In addition, the rat models pointed to sexual dimorphism, with lower severity of phenotypes in rat females compared to male mutants. This phenomenon is also observed in humans. We are convinced that the two rat models will be key to further investigating social behaviour and understanding the brain mechanisms and specific brain regions that are key to controlling social behaviour.

## INTRODUCTION

The 16p11.2 locus is a pericentromeric region found in chromosome 16, one of the most gene-rich chromosomes in our genome, for which 10% of its sequence consists of segmental duplications [1]. These elements give strong instability and induce the appearance of copy number variations (CNV) because of the recurrent non-allelic homologous recombination mechanism [2]. The most prevalent rearrangement, deletion and duplication are generated between two low copy repeats (LCR), named BP4 and BP5, and encompasses 600 kb. 16p11.2 CNVs are an important risk factor for neurodevelopmental disorders [3], including intellectual disability (ID) [4] and autism spectrum disorder (ASD) [5-9]. In addition, the deletion and duplication of 16p11.2 have been linked to epilepsy [10-12] and attention deficit hyperactivity disorder (ADHD) [13], whereas only the duplication has been related to schizophrenia, bipolar disorder and depression [14-17].

Besides, these chromosomal rearrangements have been linked to mirrored physical phenotypic effects. The 16p11.2 deletion has been associated with the risk of diabetes-independent morbid obesity and large head circumference, while the 16p11.2 duplication has been associated with low body mass index (BMI) and small head circumference [12, 18-20]. Considering this reciprocal impact on BMI and head size, it has been suggested that changes in gene transcript levels could be responsible for the symptoms associated with these CNVs. More importantly, the severity of the developmental delay and other comorbidities vary significantly in the human population with some people having an ASD or IQ below 70 and others just below average [21-23].

Animal models have been developed and characterized to investigate the interplay between genes and proteins, the consequences on brain activity and behaviour and the understanding of neurocognitive processes affected in humans. Genes of the 16p11.2 region are highly conserved on mouse chromosome 7 and several mouse models for deletion or duplication of the 16p11.2 homologous region have been generated [23-26]. Among them, our novel 16p11.2 CNV mouse models in pure C57BL/6N genetic background named *Del(7Sult1a1-Spn)6Ya*h (noted Del/+) and *Dp(7Sult1a1-Spn)6Yah* (noted Dup/+) and investigated them focusing on behaviour and metabolism [26]. We found that *Sult1a1-Spn* CNVs affect growth, weight, adiposity, activity and memory in opposite ways. Mice carrying the deletion showed weight and adipogenesis deficits, hyperactivity with increased stereotypic behaviour and novel object memory impairments. Instead, mice carrying the duplication showed weight and adipogenesis increase, hypo-activity and memory improvements. We also found that the genetic background can favour the social interaction deficits in the deletion mice model. Altogether this observation suggests that this deficit could be the consequence of the genetic context.

To generate a model presenting more suitable autistic traits, we engineered the deletion or duplication of the human homologous region 16p11.2 in the rat in two different outbred genetic backgrounds. As a model of human disease, the rat is a more sociable animal than the mouse with a large spectrum of similar and complementary behavioural assessments and the outbred genetic background, although representing a challenge, could be of interest to detect the most robust phenotypes. Rats have shown differences from mice in several models of human disease in which genetically engineered animals for the same genes have been generated [27]. The main contribution of our research is the establishment and validation of two new 16p11.2 rat models that can be helpful to test novel pre-clinical pharmacological therapies targeting specific phenotypes and finally help to improve the lives of patients.

## MATERIALS AND METHODS

### Rat lines and genotyping

The 16p11.2 rearrangement, deletion and duplication, were studied in rat models engineered through CRISPR / Cas9 technology [28] as detailed in the supplementary information. Rat models were then bred and maintained in our animal facility which is accredited by the French Ministry for Superior Education and Research and the French Ministry of Agriculture (agreement #A67-218-37) following the Directive of the European Parliament: 2010/63/EU, revising/replacing Directive 86/609/EEC and the French Law (Decree n° 2013-118 01 and its supporting annexes entered into legislation on 01 February 2013) relative with the protection of animals used in scientific experimentation. All animal experiments were approved by the local ethical committees (Approval Committee: Com’Eth N°17 and French Ministry for Superior Education and Research (MESR) with approval licenses: internal numbers 2012-009 & 2014-024, and MESR: APAFIS#4789-2016040511578546) and supervised in compliance with the European Community guidelines for laboratory animal care and use and every effort was made to minimize the number of animals used and their suffering.

### Behavioural analysis

To decipher more in detail alterations of specific cognitive functions and autistic traits in 16p11.2 CNVs rat models on two genetic backgrounds, Sprague-Dawley (SD) and LongEvans (LE), we evaluated several phenotypes with a validated rat phenotyping behavioural pipeline. We defined the protocol with tasks in which 16p11.2 mouse models showed robust phenotypes: alterations of exploration activity, object location and novel object recognition (NOR) memory, and social interaction. For SD 16p11.2 rat models, littermate animals from different crosses with four genotypes were used: wt, Del/+, Dup/+ and pseudo-disomic Del/Dup (Figure 1C). For LE 16p11.2 rat model, we used littermate animals from the wt and Del/+ cross to get mutant and control genotypes as littermates.

**Figure 1.**
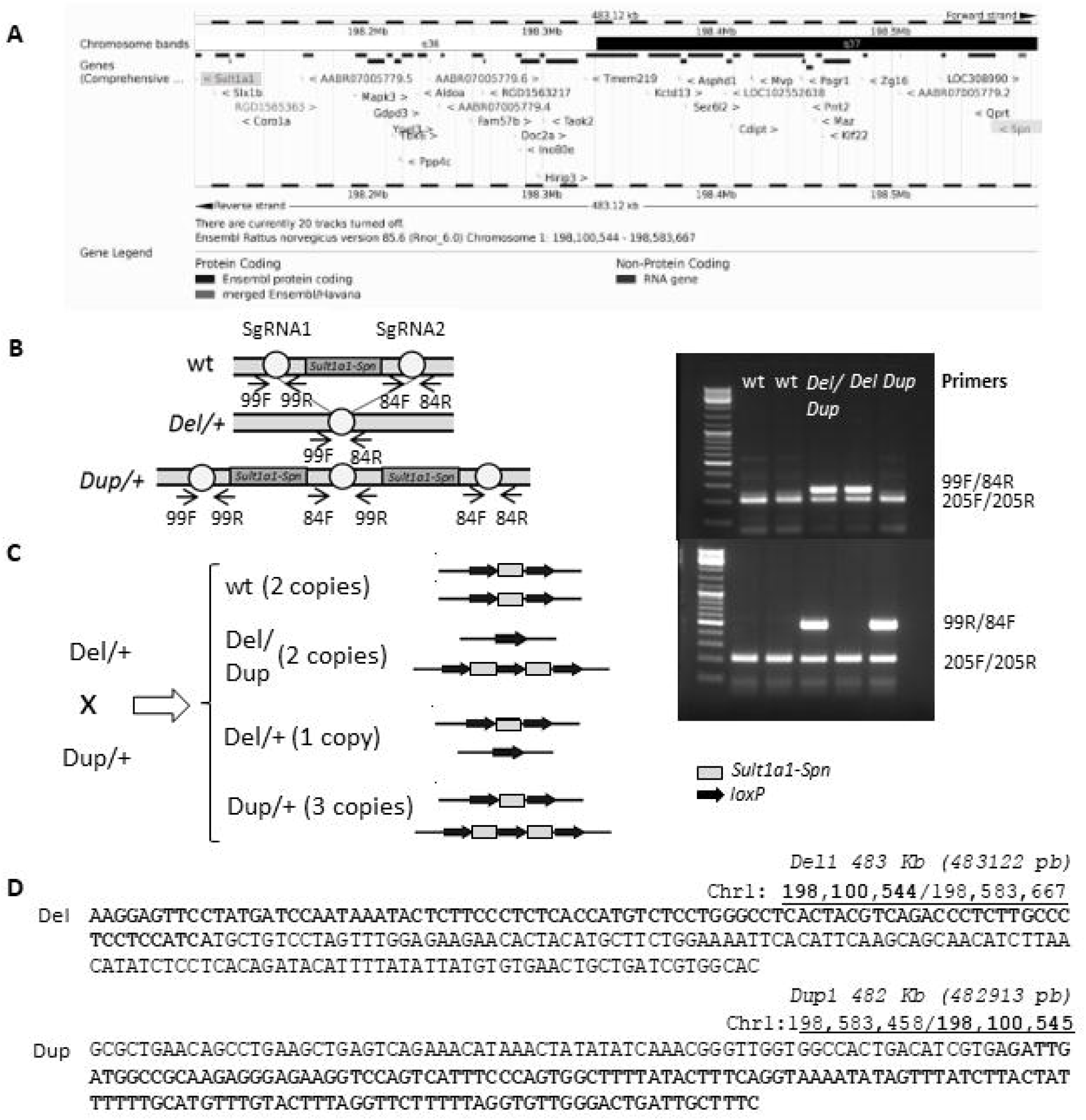
New rat SD model for the 16p11.2 syndromes. (A) The syntenic region 16p11.2 BP4-BP5 in the rat chromosome 1. (B) Left. Mutation strategy using CRISPR / Cas9 technology from the *in vitro* genome editing inside of a fertilized embryo and subsequent injection into a pseudo-pregnant female. We obtained individuals carrying the deletion and duplication of the *Sult1a1-Spn* region. Right. Molecular validation. PCR-specific products for the wt (205 bp), *Del/+* (290 bp) and *Dup/+* (500 bp) alleles. (C) Breeding strategy for obtaining wt, *Del/+, Dup/+* and *Del/Dup* littermates. (D) Junctions positions and details of the mutated genetic sequences for *Del* and *Dup* rat models. All genomic positions are given according to the UCSC rat genome browser (RGSC 6.0/rn6).

Behavioural studies were conducted in 14 to 16-week-old SD rats of both sexes separately, from 8 cohorts. Whereas LE rats were analysed separately between 19 and 24 weeks old for both sexes, from 1 large cohort. Animals were housed in couples of 2 individuals per cage (Innocage Rat cages; 909 cm^2^ of floor space; Innovive, San Diego, USA), where they had free access to water and autoclaved food (D03, Safe Diets, France). The temperature was maintained at 23±1°C and the light cycle was controlled as 12 hours light and 12 hours dark (light on at 7 am).

On the testing days, animals were transferred to the experimental room antechambers 30 min before the start of the experiments. The body weight of the animals from the SD 16p11.2 rat models was recorded at 13 weeks old whereas the body weight of animals from LE 16p11.2 rat models was recorded at 2 weeks old, and followed at weaning and 19 weeks. All tests were scored blind to the genotype as recommended by the ARRIVE guidelines [29, 30]. The protocols for open field, object location, object recognition memories and social interaction are described in the supplementary information.

### Statistical analysis

The statistical analysis of our results was carried out using standard statistical procedures operated by SigmaPlot software (Systat Software, San Jose, USA). All outliers were identified using Grubbs’ test from calculator GraphPad (GraphPad Software, San Diego) or ROUT method with a Q value of 1% from GraphPad Prism 7.01 (Motulsky et al., 2006) (GraphPad Software, San Diego) when data with nonlinear regression. Acquired data from the behavioural characterization of 16p11.2 rat models were analysed using one-way ANOVA followed by Student’s t-test and Tukey’s posthoc test whenever data presented normal distribution and equal variance. Otherwise, we used the non-parametric Kruskal-Wallis one-way analysis of variance and the Mann-Whitney U test. One sample t-test was used also to compare recognition index values to the set chance level (50%). The data to evaluate the mutant allele transmission was analysed by a Person’s chi-squared test. Data are represented as the mean ± SEM and the statistically significant threshold was p < 0.05.

### Transcriptomic analysis

Hippocampus from 4 Del/+ (noted as Del(16p11)), 5 Dup/+ (noted as Dup(16p11)) and 4 Del/Dup (noted as Del/Dup(16p11)) SD rats and 6 wt littermates for each (17weeks old), were isolated and flash frozen in liquid nitrogen after the behavioural analysis. Total RNA was prepared using an RNA extraction kit (Qiagen, Venlo, Netherlands) according to the manufacturer’s instructions. Samples quality was checked using an Agilent 2100 Bioanalyzer (Agilent Technologies, Santa Clara, California, USA). All the procedures and the analysis are detailed in the supplementary information. For the LE rat models, hippocampus from adult individuals, Del/+ and wt, male and female littermates (n=5 per group, 24 weeks old), were collected after the behavioural analysis. and treated with the same protocol.

The preparation of the libraries was done by using the TruSeq Stranded Total RNA Sample Preparation Guide - PN 15031048. The molecule extracted from the biological material was polyA RNA. The Whole genome expression sequencing was performed by the platform using Illumina Hiseq 4000 and generating single-end RNA-Seq reads of 50 bps length. The raw sequenced reads were aligned by Hisat2 against the Rno6.v96. 32623 ENSEMBL Gene Ids were quantified aligning to the Rno6.v96 assembly. HTSeq-count was used to generate the raw counts. The downstream analyses were carried on with in-house bash scripts and R version 3.6 scripts using FCROS[31] and DESeq2[32] packages to identify the DEGs. Raw reads and normalized counts have been deposited at the GEO resource database (Accession No. GSE224935 for the SD dataset and GSE for the LE dataset).

We performed the functional differential analysis [33] and grouped all the pathways into 25 functional categories (noted meta-pathways). Then, to assess the gene connectivity we build a minimum fully connected protein-protein interaction (PPI) network (noted MinPPINet) of genes known to be involved in all meta-pathways we defined as they were associated with each pathway via GO [34] and KEGG databases [35] and added regulatory information to build the final 16p11 dosage sensitive RegPPINet. We used the betweenness centrality analysis to identify hubs, and keys for maintaining the network communication flow.

### ddPCR analysis

Analyses were performed by Droplet Digital™ PCR (ddPCR™) technology. All experiments were performed following the previously published protocol [36]. Primers are described in Table S1.

Identification of central genes linked to the behavioural phenotypes

To further study the genotype-phenotype relationship in those models we combined the behavioural results and the RNA-Seq data to identify central genes altered in the models linked to the observed phenotypes using the genotype-phenotype databases GO, KEGG and DisGeNET. For this, we combined the knowledge from the human disease database DIsGeNET and the GO genesets (see Supplementary information). Then we queried our RNA-Seq data for those genes to identify those found deregulated in the datasets. Table 1 summarized the results with the genes annotated with the expression level, regulation sense on each model, log2FC and standard deviation of the log2FC.

**Table 1:**
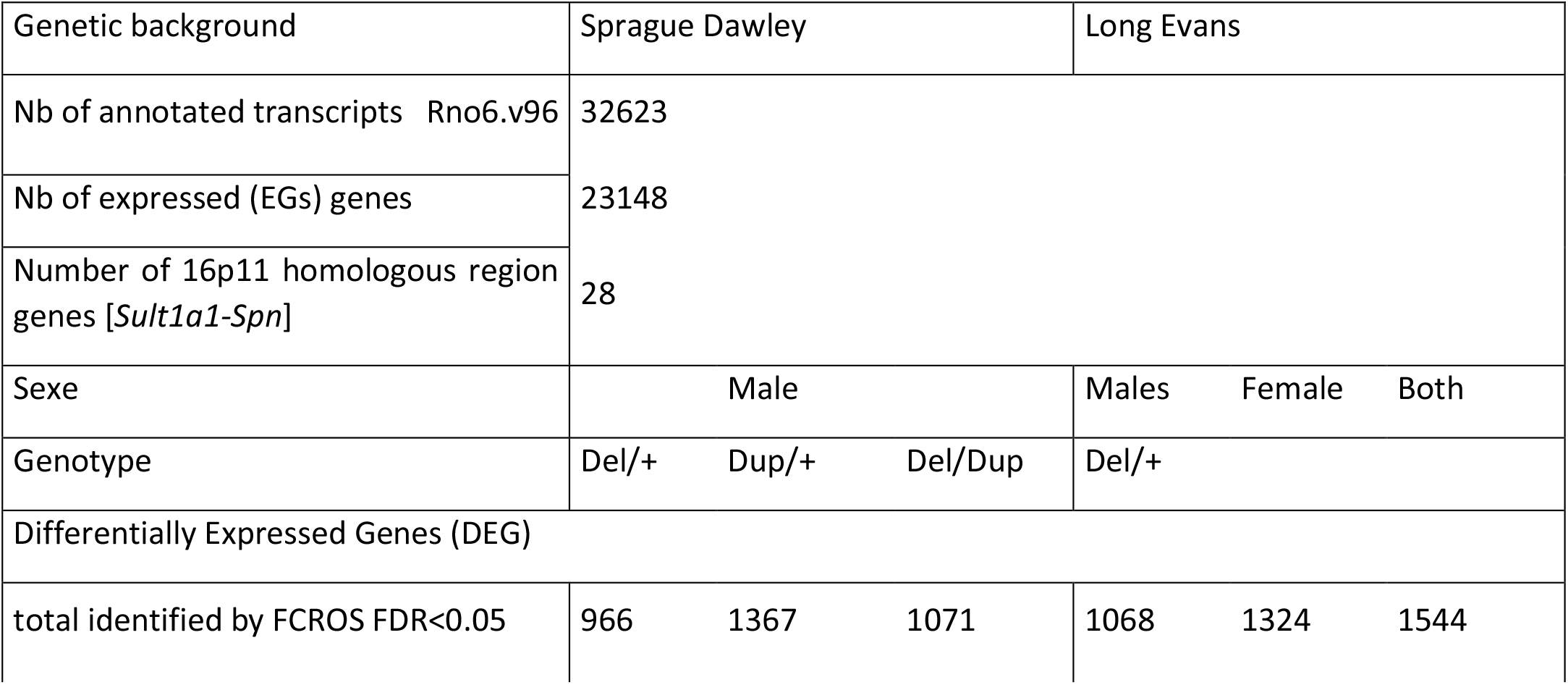
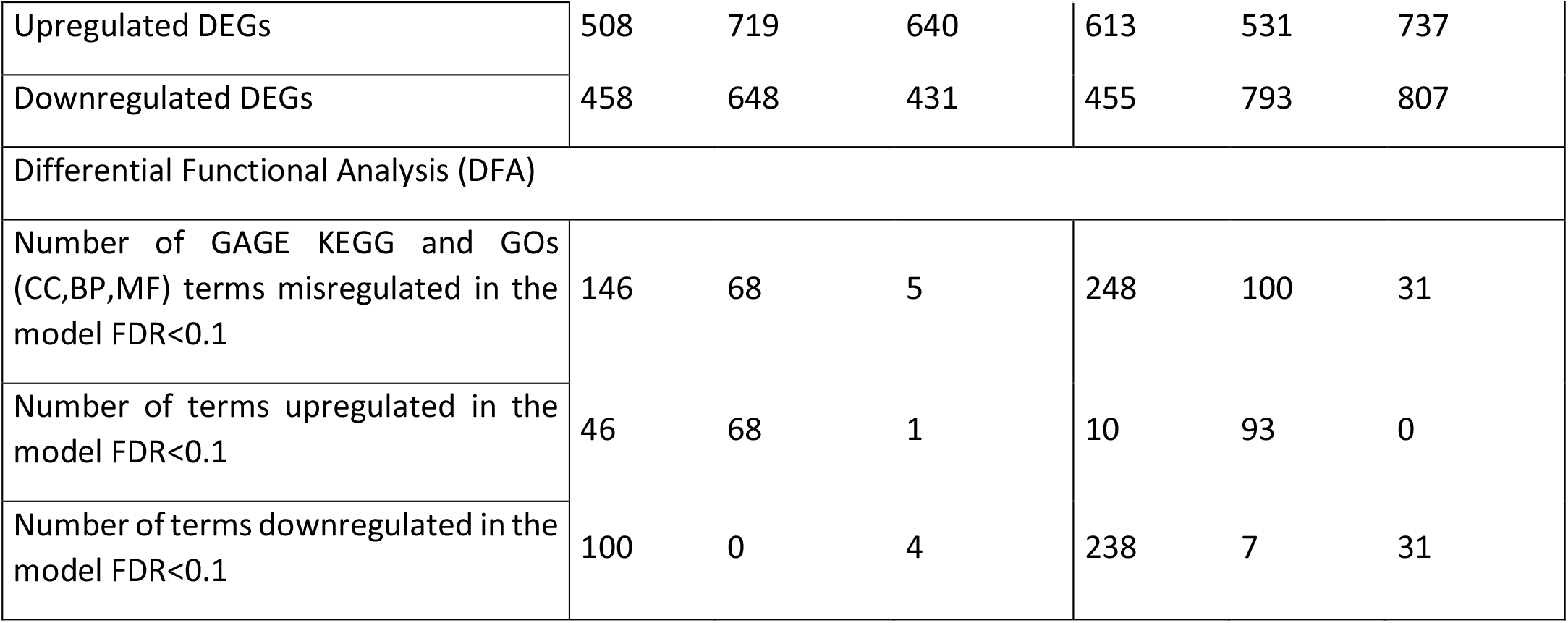
Summary of the transcriptional analysis of the SD and LE 16p11.2 models.

### Proteomic analysis

Fresh entire hippocampal tissues were isolated by CO2 inhalation/dissection of naive rats and snap frozen. Then, lysed in ice-cold sonication buffer supplemented with Complete^™^ Protease Inhibitor Cocktail (Roche).

Individual samples were disaggregated and centrifuged at 4°C for 30 minutes at 14000 rpm. Protein mixtures were TCA (Trichloroacetic acid /Acetone)-precipitated overnight at 4°C. Samples were then centrifuged at 14000 rpm for 30 minutes at 4°C. Pellets were washed twice with 1 mL cold acetone and centrifuged at 14000 rpm for 10 minutes at 4°C. Washed pellets were then urea-denatured with 8 M urea in Tris-HCl 0.1 mM, reduced with 5 mM TCEP for 30 minutes, and then alkylated with 10 mM iodoacetamide for 30 minutes in the dark. Both reduction and alkylation were performed at room temperature and under agitation (850 rpm). Double digestion was performed with endoproteinase Lys-C (Wako) at a ratio of 1/100 (enzyme/proteins) in 8 M urea for 4h, followed by overnight modified trypsin digestion (Promega) at a ratio of 1/100 (enzyme/proteins) in 2 M urea. Both Lys-C and Trypsin digestions were performed at 37°C. Peptide mixtures were then desalted on a C18 spin-column and dried on Speed-Vacuum before LC-MS/MS analysis (see Supplementary information).

## RESULTS

### General and behavioural characterization of the SD 16p11.2 rat models

The 16p11.2 region is conserved in the rat genome on chromosome 1 (Fig. 1A). Using CRISPR/Cas9 we generated first the deletion and the duplication of the conserved interval containing *Sult1a1* and *Spn1* (Fig. 1B-C) and we determined the precise sequence of the new borders of the deletion and duplication (Fig. 1D).

To carry out the behavioural analysis of the SD 16p11.2 rat models we combined the deletion (*Del/+*) and the duplication (Dup/+) for the generation of 4 groups of genotypes: wt control, Del/Dup pseudo-disomic for the 16p11.2 conserved region, Del/+ and Dup/+ littermates (Fig. 1C). Before testing, we checked the transmission of the alleles and we did not find any deviation from Mendelian rate (Table S2). Then, we analysed the effect of 16p11.2 CNVs on the weight of the 13 weeks old rats. We observed that male rats carrying 16p11.2 deletion showed a decrease in body weight compared to wt littermates, while males carrying 16p11.2 duplication did not show alterations compared to wt littermates (Table S3). The 16p11.2 rearrangements did not affect the body weight of female rats (Fig. 2A).

**Figure 2.**
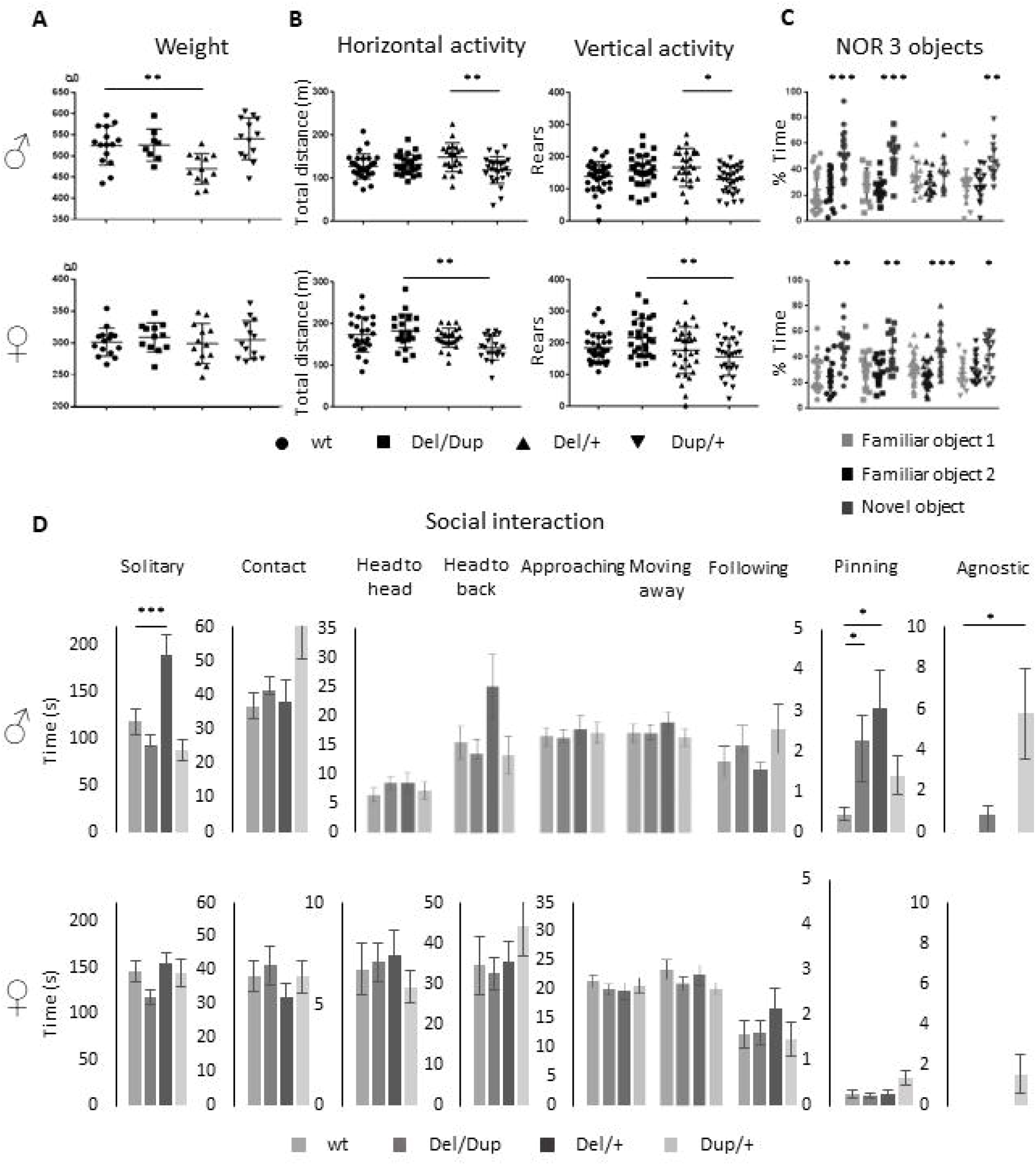
Phenotypic characterization of the 16p11.2 rat models on the SD genetic background. (A) Effects of *Sult1a1-Spn* rearrangements on body weight. Body Weight (g) of the 13 weeks old males (wt (n=15), *Del/Dup* (n=8), *Del/+* (n=12) and *Dup/+* (n=13)) and female rats (wt (n=15), *Del/Dup* (n=12), *Del/+* (n=13), and *Dup/+* (n=13)) from Del-Dup littermates. Only the deletion of 16p11.2 region caused reduced body weight in males (One way ANOVA between groups, F_(3,44)_ = 6.24; *p* = 0.001; Student t-test, *Del/+* vs wt : t_(25)_ = 3.39 *p* = 0.002). (B) Exploratory behaviour of the rat 16p11.2 models in the open field test. Male (wt (n=28), *Del/Dup* (n=26), *Del/+* (n=21) and *Dup/+* (n=27)) and female (wt (n=25), *Del/Dup* (n=22), *Del/+* (n=26) and *Dup/+* (n=22)) rats were placed in the open field for 30 min to explore the new environment. The horizontal activity was measured by the total distance travelled and vertical activity was recorded with the number of rears. Animals showed large variability and limited changes between genotypes except for *Del/+* versus *Dup/+* male (One way ANOVA between groups, Total distance F_(3,98)_ = 4.33; *p* = 0.007; Tukey’s post hoc tests : *Del/+* vs. *Dup/+* : *p* = 0.004; Rears : F_(3,117)_ = 3.55; *p* = 0.017; Tukey’s post hoc tests: *Del/+* vs. *Dup/+*: *p* = 0.016) and *Del/Dup* versus *Dup/+* female (Kruskal-Wallis one-way analysis of variance, Total distance : H_(3)_ = 14.18; *p* = 0.003, Mann-Whitney Test: *Del/Dup* vs. *Dup/+* : *p* = 0.002); (One way ANOVA between groups, Rears : F_(3,116)_ = 4.83; *p* = 0.003; Tukey’s post hoc tests: *Del/Dup* vs. *Dup/+*: *p* = 0.002). (C) Novel object recognition memory task of the rat 16p11.2 models after 3 hours of retention with 3 objects. Male rats from different genotypes (wt (n=26), *Del/Dup* (n=16), *Del/+* (n=14) and *Dup/+* (n=16)) and female rats (wt (n=18), *Del/Dup* (n=16), *Del/+* (n=26), and *Dup/+* (n=17)) were tested for the novelty recognition. The graphs show the percentage of time spent by the animals exploring a novel object compared to the time spent exploring two familiar objects. We compared the recognition index, like the percentage of exploration time of the new object, to the level of chance (33.3 %). Only the *Del/+* males showed impairment in the recognition index (One sample t-test: wt (t_(25)_ = 4.6; *p* = 0.0001), *Del/Dup* (t_(15)_ = 2.82; *p* = 0.01), *Del/+* (t_(13)_ = 0.34; *p* = 0.74) and *Dup/+* (t_(14)_ = 3.58; *p* = 0.0030) compared to all the other genotypes in males and females. Surprisingly, no change was observed in the *Del/+* females (One sample t test: wt (t_(17)_ = 3.67; *p* = 0.002), *Del/Dup* (t_(16)_ = 3.08; *p* = 0.01), *Del/+* (t_(25)_ = 3.83; *p* = 0.0008) and *Dup/+* (t_(16)_ = 2.3; *p* = 0.035). (D) Social interaction of the 16p11.2 rat models. Male (wt (n=15), *Del/Dup* (n=14), 16p11.2 *Del/+* (n=10) and *Dup/+* (n=14)) and female (wt (n=14), 16p11.2 *Del/+* (n=15), 16p11.2 *Dup/+* (n=12) and *Del/Dup* (n=15)) rats were tested for impairment of social interaction in pairs of individuals from different home cages with the same genotype. The *Del/+* male rat showed increased solitary time (One way ANOVA between groups, Solitary behaviour: F_(3,49)_ = 9.85; *p* < 0.001; Tukey’s post hoc tests : *De/+* vs. wt : *p* < 0.001, *Del/+* vs. *Del/Dup* : *p* < 0.001 and *Del/+* vs. *Dup/+* : *p* = 0.005). and pinning behaviour with *Del/Dup* (Kruskal-Wallis one-way analysis of variance: H_(3)_ = 8.66; *p* = 0.03; Mann-Witney test: *Del/+* vs. wt: *p* = 0.04; *Del/Dup* vs. wt: *p* = 0.01) while Dup/+ males are more agnostic (Kruskal-Wallis one-way analysis of variance, H_(3)_ = 13.63; *p* = 0.003; Mann-Witney test: *Dup/+* vs. wt: *p* = 0.01; *Dup/+* vs. *Del/+*: *p* = 0.02). No altered social behaviour has been detected in females. (* *p* < 0.05; ** *p* < 0.01; *** *p* < 0.001).

As a first analysis for consequences of 16p11.2 CNVs in neuronal function, we measured spontaneous locomotion activity and exploratory behaviour in the open field test (Table S4). The horizontal activity was measured through total travelled distance, whereas the vertical activity was analysed by the number of rears. As shown in Fig. 2B, increased variability was observed in the distance travelled and the rearing activity. The only significant differences were found between extreme genotype male Del/+ versus Dup/+ and between female Del/Dup versus Dup/+ for both the distance travelled and the rearing activity in the open field.

Then, we carried out the novel object location recognition test and the novel object recognition test, common assays for assessing impaired memory in rodents. For the novel object location recognition, animals were challenged to discriminate a moved object from an unmoved object. We first evaluated the performances of males and females separately, and as no significant sex differences were noted, we combined these data across both sexes. No difference was observed between genotypes in the retention session (Kruskal-Wallis one-way analysis of variance: H_(3)_ = 5.61; *p* = 0.132; Fig. S1A). We also compared the recognition index of the animals, i.e. the percentage of exploration time of the new object location, with the level of chance (50%). The new object position was always explored more than the object not moved for all the genotypes.

Following these observations, we next assessed novel object recognition from a first paradigm, based on the protocol used for the CNVs 16p11.2 *Sult1a1-Spn* mouse model. The animals should be able to differentiate an object observed previously during the acquisition phase from a novel object presented during the retention phase (Fig. S1B). All four genotypes engaged in similar levels of novelty discrimination (One way ANOVA: F_(3,88)_ = 0.038; *p* = 0.99). We also compared the recognition index with the level of chance (50%). A general preference was observed for the new object compared to the familiar object. The rat 16p11.2 models displayed correct recognition memory, with an increased time of the new object exploration spent by rats compared to mice [26].

Thus, we used a more complex object recognition paradigm with 3 different objects (Fig S1C). In this case, the male Del/+ carriers showed impairment in the discrimination of the novel object compared to all the other genotypes (Fig. 2C). Nevertheless, no alteration was observed in the females, with all the genotypes able to discriminate the novel versus the two familiar objects.

Finally, the last task focused on studying rat social interactions by analysing different social behaviour [37] (Fig. 2D). The Del/+ male displayed significantly increased time in solitary compared to all genotypes. In addition, 16p11.2 Del/+ was associated with the presence of more pinning, a behaviour to exert dominance. Interestingly the pinning behaviour was found also in the Del/Dup male animals. Furthermore, male Dup/+ showed increased agnostic behaviour compared to all genotypes. Surprisingly we did not see any social phenotypes in females of the Del/+ or Dup/+ genotype.

### Behavioural characterization of the LE *Del/+* 16p11.2 rat model

Elucidating the genetic mechanisms by which some CNVs influence neurodevelopment requires a rigorous quantitative analysis of the human phenotype but also the establishment of validated model systems in which the phenotypic diversity is conserved. Based on previous results obtained on a pure inbreed mouse C57BL/6JN, or on a mixed B6.C3H genetic background [26] and now on a rat outbred SD model, we decided to investigate if the phenotypes were robust enough, and the deficits preserved in a different background. In addition, we considered it pertinent to verify if the females were equally more resilient to the deletion of the 16p11.2 region than the males in this new genetic background. For this purpose, we engineered the deletion of the homologous region to the human 16p11.2 BP4-BP5 locus in a rat LE outbred strain and we verified the location of the specific Del interval (Fig. S2). Interestingly the transmission of the Del/+ allele was affected in females of the LE 16p11.2 model (Table S1) while males did not show any significant change. We also evaluated the body weight of this new model for both sexes and found that the deletion of the 16p11.2 region caused a decrease in the body weight on the LE genetic background in both sexes (Fig. 3A; Table S5). Then, we proceeded to study the behaviour using the same battery of behavioural tests (Table S5). We analysed the locomotion and exploratory activity of our second model. First, we observed a reduction in the variability between the data of each individual regarding the results observed in the SD model. In addition, we detected a significant increase in horizontal activity among individuals carrying the 16p11.2 region deletion for both sexes. Finally, the Open Field test showed a significant increase in vertical activity, evaluated as the total number of rears, only in male rats. These results translate into the presence of stereotypical behaviours in this model associated with genotype and sex (Fig. 3B).

**Figure 3.**
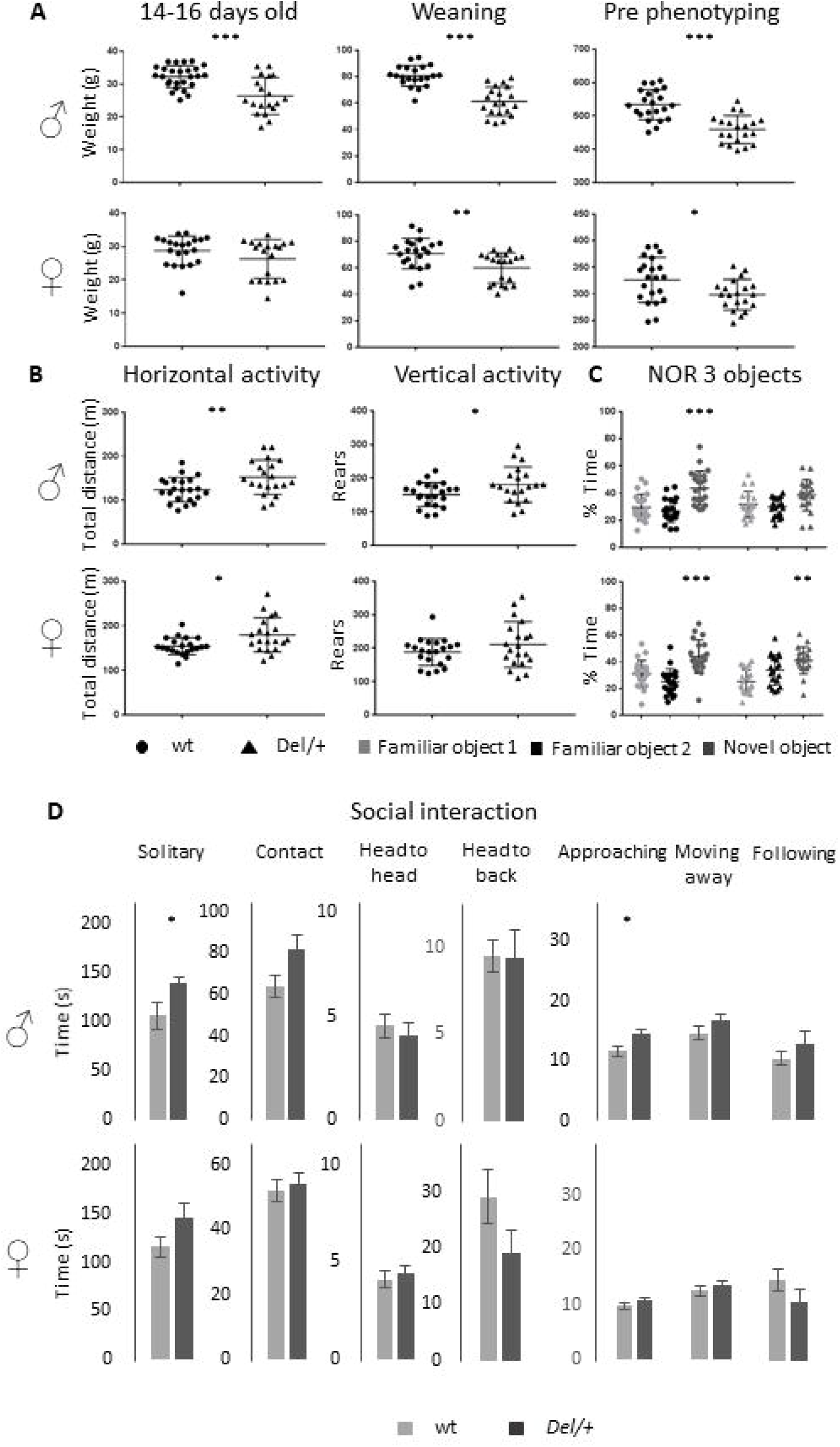
Phenotypic characterization of the 16p11.2 rat models on the LE genetic background. (A) Effects of *Sult1a1-Spn* deletion on body weight of LE 16p11.2 *Del/+* rat model. Left. Body weight (g) of the 14-16 days old males (wt (n=27) and *Del/+* (n=19)) and female rats (wt (n=21) and Del/+ (n=19)). Our observations showed a decreased body weight in *Del/+* males compared to wt littermates (Student’s t-test : t_(44)_ = 3,819; *p* < 0,001). Central. Body Weight at weaning of the males (wt (n=21) and *Del/+* (n = 20)) and female rats (wt (n = 21) and *Del/+* (n=20)) from LE *Del/+* littermates. The deletion of the 16p11.2 region caused body weight decrease in male (Student’s t-test : t_(39)_ = 6,550; *p* < 0,001) and female individuals (Student’s t-test : t_(39)_ = 3,036; *p* = 0,004) compared to wt littermates. Right. Body weight of 16p11.2 *Del/+* male (wt (n=22) and *Del/+* (n = 20)) and female (wt (n=21) and *Del/+* (n = 20)) littermates during the first week of phenotype analysis. The male and female individuals carrying the deletion of the interest region continued to show a decrease in their body weight throughout their development (Student’s t-test for males : t_(40)_ = 5,550; *p* < 0,001; Student’s t-test for females : t_(39)_ = 2,451; *p* = 0,019). (B) Open Field test results illustrate the exploratory activity of the LE 16p11.2 rat model. Male (wt (n=22) and *Del/+* (n=20)) and female (wt (n=21) and *Del/+* (n=20)) littermates were analyzed for horizontal and vertical activity. The 16p11.2 deletion caused increased horizontal activity in our model regardless of the sex of the animals (Student’s t-test for males : t_(40)_ = -2,726; *p* = 0,009; Mann-Whitney U Statistic for females : T = 510,000; *p* = 0,02). However, the deletion of one copy of the interest region caused increased vertical activity only in male individuals (Student’s t-test : t_(40)_ = -2,174; *p* = 0,036). (C) The deletion of the 16p11.2 region causes a novel object recognition memory disorder in our rat model on LE genetic background. For the NOR test with 3 objects, the recognition index reflects the ability of rats to recognize the new object from the 2 familiar objects after a 3 h delay. Males mutant animals (wt (n= 22) and *Del/+* (n= 20)) were impaired to recognize the new object when we compared the recognition index, like the percentage of exploration time of the new object, to the level of chance (33.3 %) (One sample t-test: wt (t_(21)_ = 3.94; *p* = 0.0008) and *Del/+* (t_(19)_ = 2.02; *p* = 0.0569). However, the females of this model (wt (n= 21) and *Del/+* (n= 20)) showed a preference for the new object that is reflected in a recognition index significantly higher than the level of chance (One sample t-test: wt (t_(20)_ = 3.92; *p* = 0.0008) and *Del/+* (t_(19)_ = 3.4; *p* = 0.003). (D) Evaluation of the behaviour of the LE 16p11.2 rat model in the social interaction test. The male (wt (n=11) and *Del/+* (n=10)) and female (wt (n=10) and *Del/+* (n=10)) of our second model were analysed separately from the observation of different events in pairs. The *Del/+* male rat showed increased solitary time (Student’s t-test : t_(18)_ = -2,229; *p* = 0,039) and approaching behaviour (Student’s t-test : t_(19)_ = -2,679; *p* = 0,015). No altered social behaviour has been detected in females. (* *p* < 0.05; ** *p* < 0.01; *** *p* < 0.001).

Next, we performed the object location memory test (Fig S3A, Table S6). Our objective was to confirm, as in the case of the SD model, that the deletion of a copy of the 16p11.2 region has no impact on the object location memory in our second rat model. Considering that the 3-object discrimination protocol for the NOR test was the most appropriate, we decided to test the LE model (Fig S3B and Fig. 3C). Thus, this task showed that male mutant individuals, unlike control individuals, did not show an exploration preference for the new object. Furthermore, the object recognition index of these animals is not significantly higher than 33.3%. Instead, females carrying the 16p11.2 deletion on LE background did not develop any disorder in object recognition memory, showing a recognition index higher than the 33.3% chance level.

To get an animal model more relevant to autism with robust social behaviour phenotypes shared among genetic backgrounds, we evaluated the LE model during the social interaction test (Fig. 3D). Among the most interesting observations, we found that male mutant individuals spent significantly more time alone than control individuals. This phenotype was also observed in the SD 16p11.2 deletion model (Fig. 2D). In addition, curiously, we discovered that these animals spent more time approaching their partner and we did not detect cases of pinning or agnostic behaviour. In the case of LE females, the deletion of the 16p11.2 region had no effect on the development of social phenotypes for the evaluated events, as observed for the mutant SD females Overall the behaviour results we identified were robust, as were observed similarly in the LE background to those obtained from the SD background in the 16p11.2 Del/+ model.

### Expression analysis shed light on the pathways altered by a genetic dosage of the 16p11.2 homologous region

We investigated the gene expression profile of Del/+ and Dup/+ 16p11.2 male SD rat hippocampi by RNA-seq. After performing the DEA analysis using FCROS we identified 966 and 1367 genes dysregulated (DEGs) in Del and Dup models respectively (Tables 1). Using those DEGs, we computed a PCA to assure the profile of expression of the DEGs could cluster our samples by their different gene dosage (Fig. 4A). Additionally, to assure the quality of our data, we looked at the Euclidian distance between samples, calculated using the 28 genes susceptible of the dosage effect of the 16p11 region, and indeed all cluster by genotype (Fig. 4B). Moreover, most of the genes of the region are following the gene dosage on the models. Looking at the FC profile across the region in the duplication, we found one gene with decreased FC expression LOC102552638, and several highly expressed genes as *Gdpd3*, two rat-specific *ABR07005778*.*1, AABR07005779*.*7*, and *Zg16*. Interestingly, in the deletion, we found 2 genes with an unpredicted increased expression, one outside of the region *RF00026*, possibly due to a bordering effect and another inside the region, *Zg16*. Overall, the gene expression dysregulation corroborated this region’s genetic dosage in the different models (Fig. 4C).

**Figure 4:**
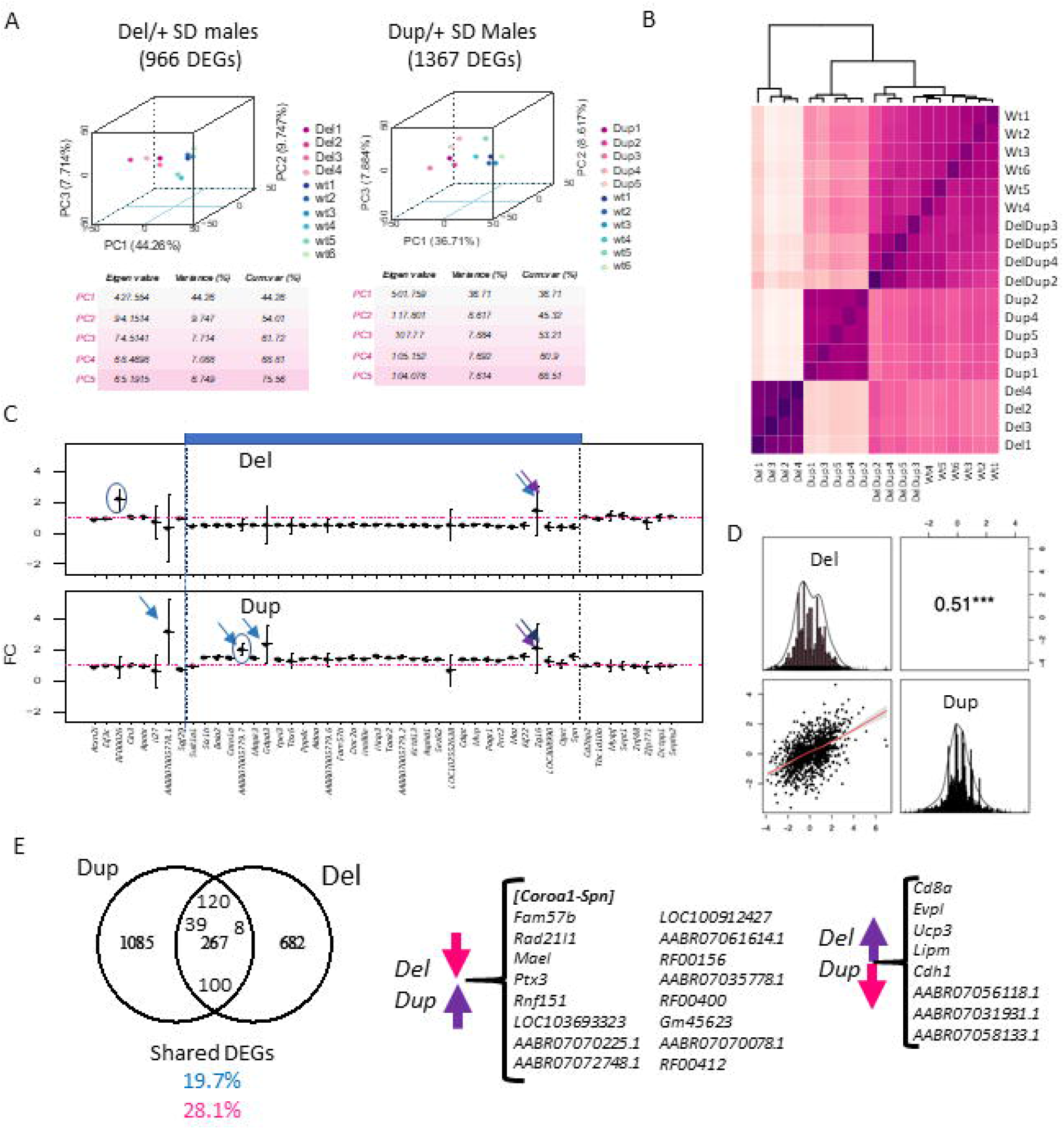
Gene expression analysis at the transcriptome and proteome level in the 16p11Del and Dup male rat hippocampi. (A) 3D principal component analysis (PCA) on the DEGs for each adult hippocampal sample allows isolating the animals carrying the 16p11.2 deletion (Del/+) and the animals with the 16p11 region (Dup/+) in comparison with the wild-type littermates. from disomic (wt) adult hippocampi. (B) Homogeneity plot showing the gene dosage effect of the 28 genes in the 16p11.2 region and how the samples cluster by Euclidian distance. (C) Fold change expression levels of the genes analysed by RNA-Seq homologous to the 16p11.2 region in rat chromosome 1 (Rno1). The genes are displayed following the order of their genomic start site coordinates. The 16p11.2 interval is indicated in blue. (D) Distribution and correlation diagram showing the DEGs in common between Del and Dup models and the model-specific DEGs. The shared DEGs correspond to 19.7% and 28.1 % of the total DEGs identified respectively for the Dup and Del models. (E) Venn diagram showing in the left panel, the DEGs in common between the SD Del and Dup models and SD male transcriptomics datasets and the background-specific DEGs. Highlighted inside the common genes those upregulated in both models, 120, downregulated in both 100, and regulated in opposing regulatory sense between Dup and Del models (39 upregulated in Dup and 8 downregulated in Dup).

Looking at the expression of DEGs in both models, only 51% were strongly correlated to gene dosage (Fig. 4D; Table S7). Many genes of the 16p11 region (*AldoA, Mapk3, Cdipt, Coroa1, Kctd13, Ino80e, Mvp, Slx1b* and *Ppp4c*) showed a level of expression in RNA-Seq following a gene dosage effect that was confirmed by ddPCR (Fig. S4). Then, we wondered whose genome-wide DEG expression levels were positively, or negatively, correlated with the gene dosage. To answer this question we fit a linear model considering CNVs as follows (lm(log2FC ∼ CNV) == y ∼ b0 + b1*CNV)). We found six genes of the region following a positive gene dosage effect *Aldoa, Sez6l2, Bola2, Kif22, Rad21l1, Ptx3, and Mael* with another gene, named *Chad*, out of the region, and presenting a negative correlation in the dosage model (Table S7). Overall, 267 DEGs were commonly dysregulated in the Del/+ and Dup/+ models (19.7 % shared DEGs of the total Dup/+ DEGs or 28.1% of the total Del/+ DEGs). Of those common 267 DEGS between models, 100 DEGs were downregulated in both models and 120 upregulated in both. Therefore, some functionalities should be commonly altered independently of the dosage. Nevertheless, 39 DEGs were following the region dosage effect (upregulated in the Dup/+ and downregulated in the Del/+), including the genes on the interval *Coroa1-Spn* and others like *Fam57b, Rad21l1, Mael, Ptx3* or *Rnf151*, found elsewhere in the genome, and 47 genes were altered in opposing regulatory sense. In particular, a few DEGs were following a negative dosage correlation such as *Cd8a, Evpl, Ucp3, Lipm* and *Cdh1* being upregulated in the Del/+ and downregulated in the Dup/+ (Fig. 4E).

To go further, we performed the differential functional analysis (DFA) using gage [38]. We found 146 and 68 pathways altered in the hippocampi of Del/+ and Dup/+ models respectively. No downregulated pathway was found in the Dup/+ model whereas both up and downregulated pathways were found in the Del/+ hippocampi. After grouping the pathways inside of functionality-based defined meta-pathways [33] (Fig. 5A; Table S8). Although in the Dup/+ model, there were several groups with a higher number of upregulated pathways compared to the Del/+ model, as synaptic meta-pathway, signalling or transcription and epigenomic regulation, many downregulated pathways were observed in the Del/+ model, except the “transcription and epigenomic regulation” meta-pathway not being affected. Moreover, as expected from the DEA analysis, we indeed were able to identify 23 pathways that were commonly shared and upregulated in both Del/+ and Dup/+ (Fig. 5B) with most of them being related to morphogenesis with the primary cilium, and four others unrelated. Oppositely “synaptic and Synaptic: other pathways” and “metabolism” functions were more affected in the Del/+ condition, while “transcription and epigenomic regulation” or “hormone regulation” were more perturbed in the Dup/+ model (Fig. 5A and 5C).

**Figure 5:**
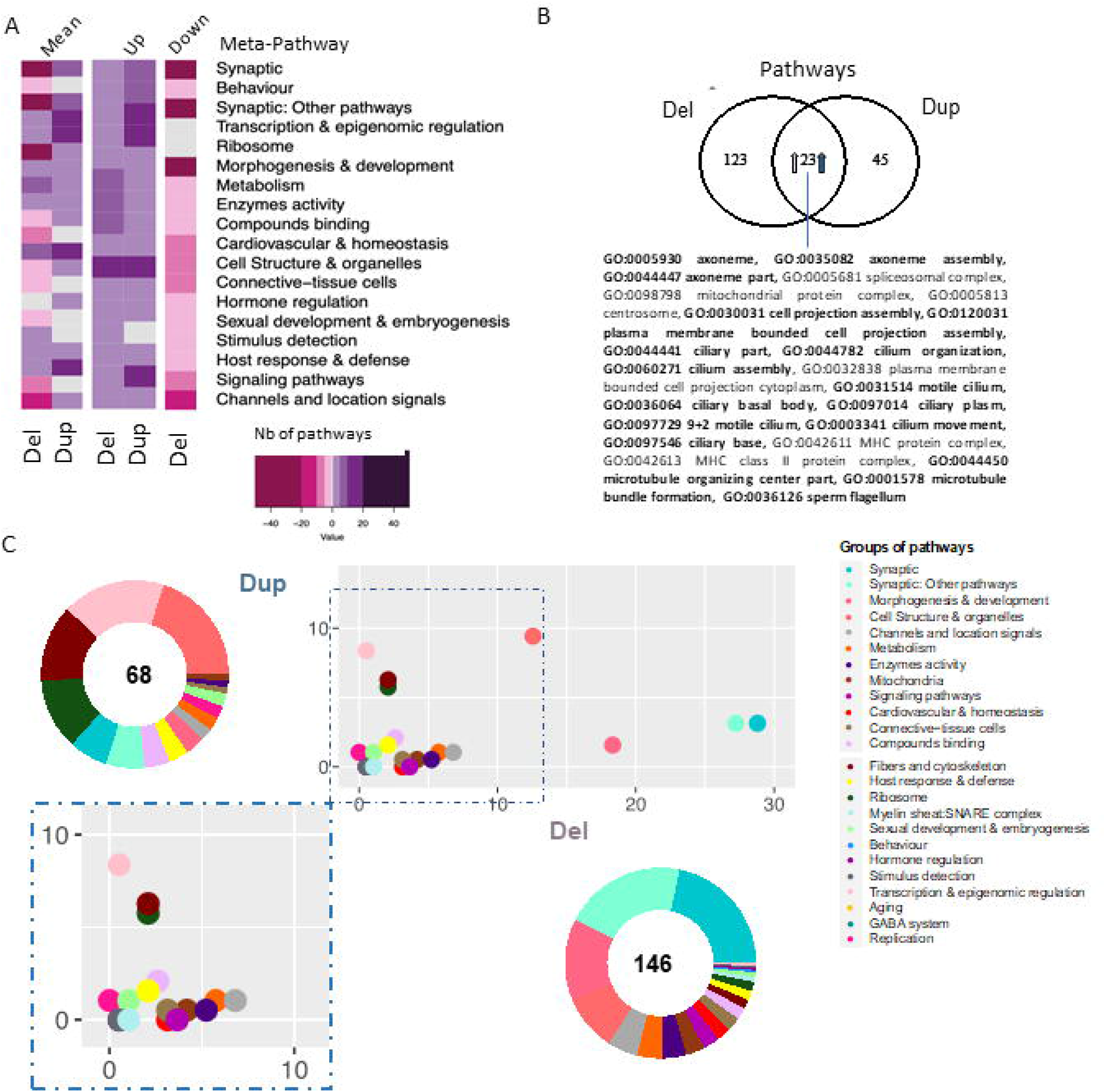
Pathway analysis of 16p11.2 SD rat Del and Dup male model based on the transcriptome of the hippocampi. (A) Heatmap representation of the number and regulation sense of the pathways of the Del and Dup models. Pathways identified using the GAGE R package and filtered by q-value cut-off < 0.1, were grouped in the meta-pathways shown on the ordinate. The colour key represents the number of pathways within the meta-pathways 50,20,10,5,0. The minus or pink colour represents down-regulated pathways, the white colour represents no pathway found in the meta-pathway and the purple or positive numbers stand for up-regulated pathways respectively. (B) Venn diagram highlighting the 23 pathways upregulated in both models and the existence of model-specific functional alteration. The percentage of shared pathways reached 33% or 15% of the total altered pathways in Dup16p11) and Del(16p11) models respectively. Seventeen of those pathways are also found dysregulated in Del/Dup (in bold). (C) Ratio plot showing the inter-model comparison of the percentage of pathways included on each meta-pathway (group of pathways) normalised by the total number of unique pathways per meta-pathway. The x-axis and y-axis represent the SD male Del and Dup data respectively. Outside a doughnut plot representing in the centre the number of total altered pathways found by gage analysis one each dataset and the percentage of pathways altered included on each meta-pathway is represented on the coronal area under each meta-pathway. The meta-pathways are defined in the accompanying legend.

### CUL3 and MAPK3 functional subnetworks are central to the 16p11 dosage susceptible regulatory protein-protein interaction network

Then, we built the rat 16p11 dosage susceptible regulatory protein-protein interaction network (RegPPINet; Fig. 6A) using as seeds all the genes identified by gage as altered in the Del/+ and/or in the Dup/+ SD models. We aimed to gain some insights into the possible molecular mechanism altered due to the gene dosage of the region. After performing the betweenness centrality analysis and analysing the topology of the most central network we identified 47 main hubs. Several of those hubs involved genes from the region (Fig. 6A) and interestingly we identified a few central genes linked to synaptic deregulation, re-enforcing the fact that synaptic dysfunction was one of the main alterations due to the dosage change of 16p11. These most important hubs in terms of betweenness were Chd1, Gli1, Plg, Coro1a, Epha8, Disc1, Spag6l, Cfap52 *or* Sema3a. However, if we consider the most connected gene, by the sole degree of regulatory interactions, then the 16p11 dosage RegPPINet pointed to MAP3 and CUL3 (Fig. 6C and 6D). The first subnetwork is centred on MAPK3 with expressed genes also found altered in both Del/+ and Dup/+ models with opposite regulatory senses (like *Cdh1* or *Mvp*). In addition, we found 3 genes on this subnetwork, *Gdnf, Gata4* and *Atp1a4* downregulated in both models and one gene *Itgb6* upregulated in both. The CUL3 network involved several genes whose expression was found altered in both Del/+ and Dup/+ models with mirroring regulatory effects for *Kctd13, Doc2a, Kif22, Rad21l1, Ppp4c* or *Asphd1*.

**Figure 6:**
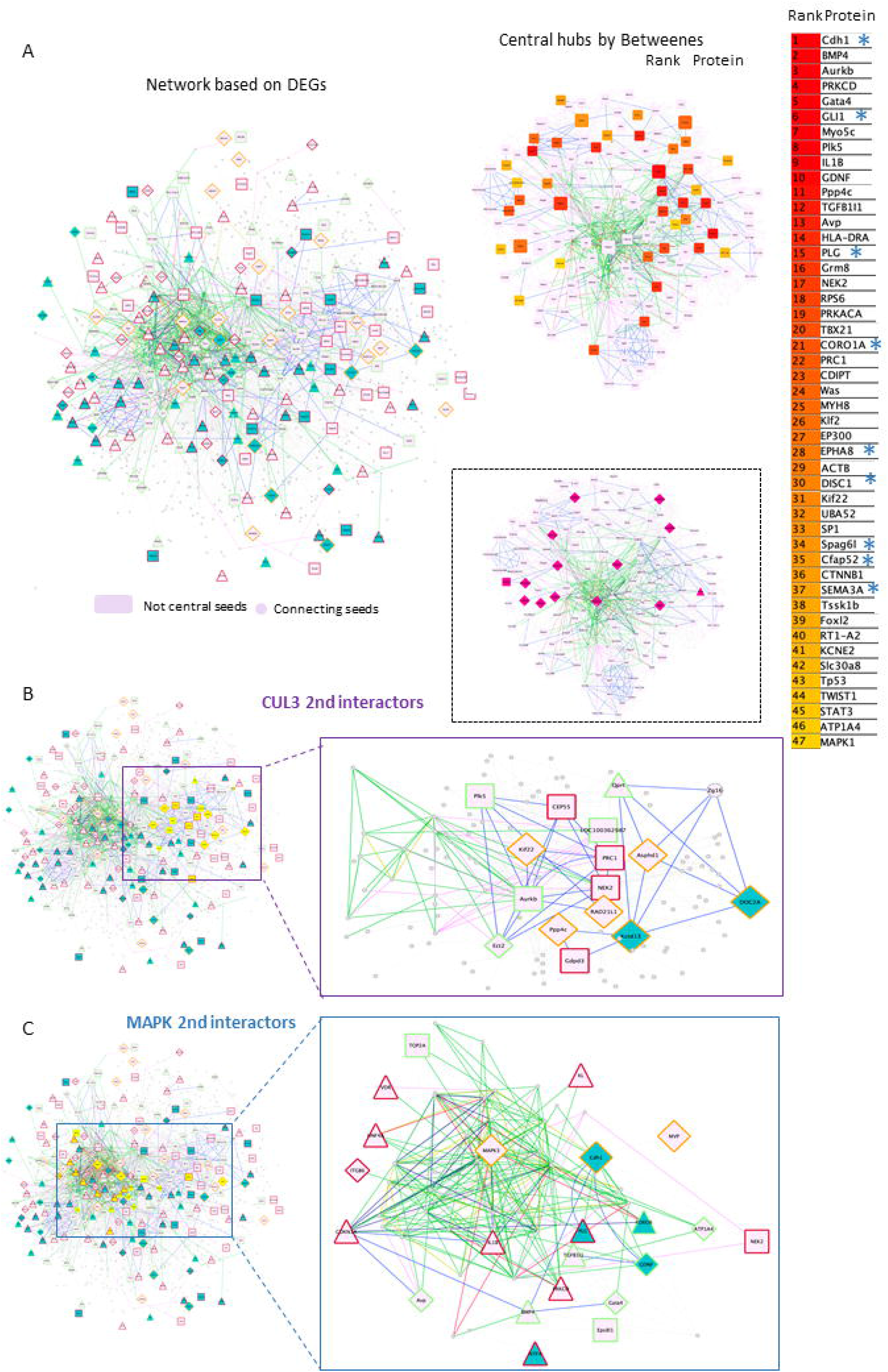
Protein-protein interaction networks altered due to the gene dosage effect based on the transcriptome analysis of the 16p11 SD rat models. (A) Left panel, a full rat protein-protein interaction network (RegPPINet) built using as seeds all the genes identified by gage as altered in the Del or Dup SD models visualized using the edge weighed spring embedded layout by betweenness index in Cytoscape. On the right panel, highlighting the main central nodes of the rat dosage susceptible RegPPINet network. On the bottom panel, highlight in pink the 16p11 region genes. The full RegPPINet was built by querying STRING and selecting the PPIs with a medium confidence score (CS=0.4) coming from all sources of evidence. The shapes of the nodes represent the following information: Shapes: i) Pallid pink ellipses: represent connecting proteins added to assure the full connectivity of the network; Then the genes identified by GAGE after q-Val <0.1 cut-offs to be contributing even slightly, to any pathway alteration and also identified in the DEA analysis by Fcros. The top 50 central genes are listed on the right side with genes known to be involved in synaptic pathways (*). (B) Second-level interactors of CUL3 were extracted from the main network. Left panel, to show in yellow all the genes that are second-level interactors of CUL3. On the right panel, the extracted subnetwork centred around CUL3 to identify the molecular regulatory mechanisms known to exist between the interacting partners. (C) We extracted from the rat RegPPINet the second-level interactors of MAPK family proteins (proteins MAPK 1,3,8;9;14). Left panel, the full RegPPINet network can be observed highlighting in yellow all the genes that are second-level interactors of CUL3. On the right panel, the extracted subnetwork centred around MAPK proteins to identify the molecular regulatory mechanisms known to exist between the interacting partners. The shapes of the nodes represent the following information: Shapes: i) Pallid pink ellipses: represent connecting proteins added to assure the full connectivity of the network; Then the genes identified by GAGE after q-Val <0.1 cut-offs to be contributing even slightly, to any pathway alteration and also identified in the DEA analysis by Fcros. Rectangles represent genes identified uniquely in Dup transcriptomes while triangles are for genes identified uniquely in Del transcriptomes and diamond shapes for genes identified as DEGs in both models. The edges colour represent the type of interaction annotated by following the PathPPI classification [50], and ReactomeFIViz annotations as follows i) The GErel edges indicating expression were coloured in blue and repression in yellow. ii) PPrel edges indicating activation were coloured in green, inhibition in red. Iii) Interactions between proteins known to be part of complexes in violet. Iv) Predicted interactions were represented in grey including the PPI interactions identified by STRING DB [51] after merging both networks. The nodes bordering colour represent If the gene was found upregulated in both models (red), downregulated in both (green) or in the mirroring regulatory sense (orange).

**Figure 7:**
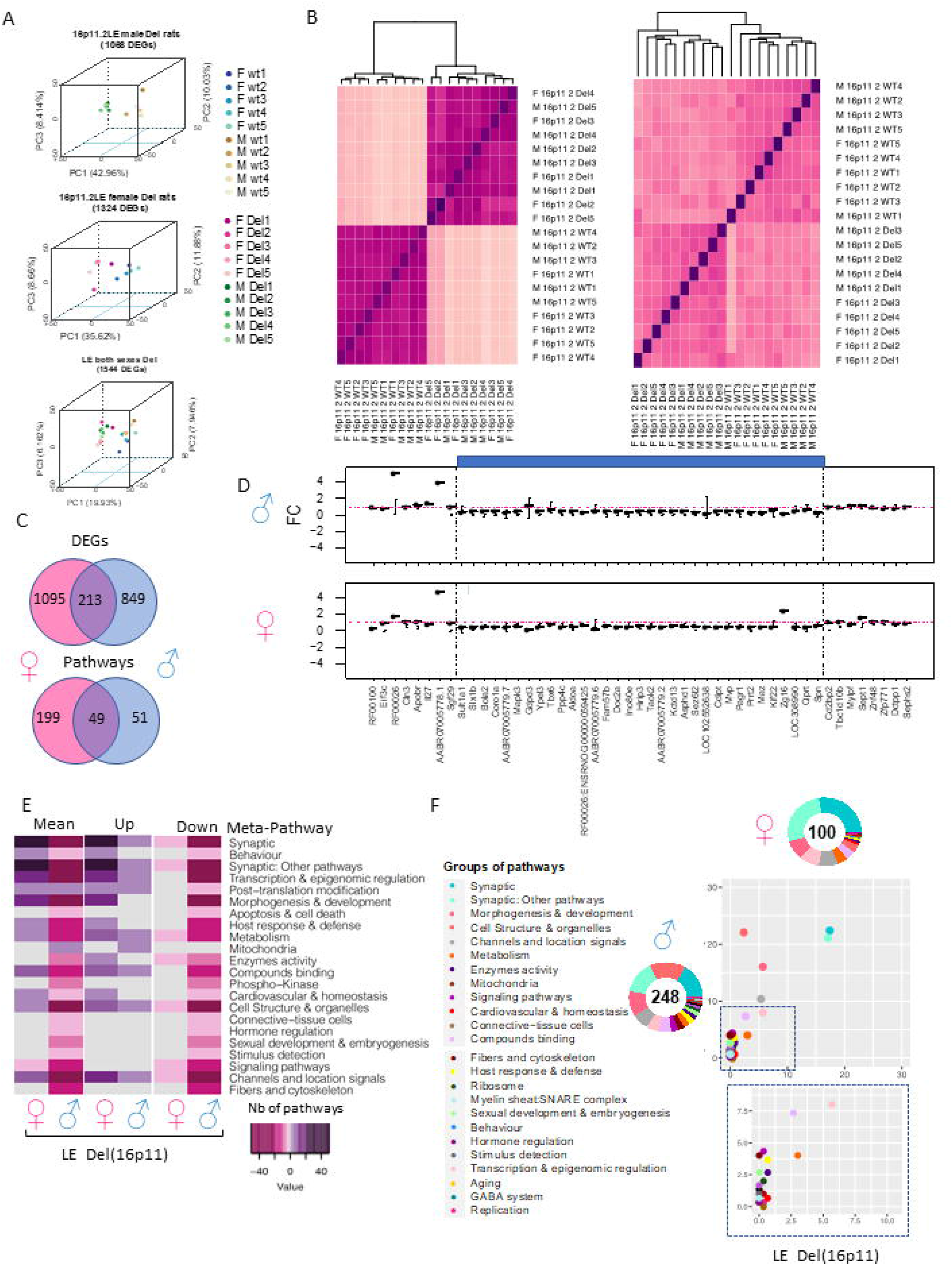
Gene expression analysis of 16p11 LE Del and control littermate (wt) male and female rats. (A) 3D-PCA on the DEGs for each adult hippocampal sample allows to isolate the LE rats carrying the 16p11.2 deletion (Del) in comparison with the wild-type littermates in both sexes as shown in the upper and middle plots. As shown in the bottom plot even though there are some differences by sex the genotype effect is the mayor difference between the animals and its variability is explained in the first component « PC1 ». (B) Homogeneity plot showing the gene dosage effect of left) the 28 genes in the 16p11.2 region and how the samples cluster by Euclidian distance, and right) all DEGs identified in both male and female datasets. (C) Venn diagram showing in the upper panel, the DEGs were found common between the male and female LE datasets. The shared DEGs correspond to 16.2% and 20 % of the total DEGs identified respectively for the female and males. In the bottom panel, Venn diagram showing the pathways in common between the male and female LE datasets. The shared pathways correspond to 19.7% and 49 % of the total pathways identified. (D) Fold change expression levels of the genes from the region homologous to 16p11.2 in Rno1. The genes are displayed following the order of their genomic start site coordinates. The deleted areas for each model appear shaded in blue. (E) Group of meta-pathways showing up or down-regulation with a colour key corresponding to the number of pathways within the meta-pathways. (F) Ratio plot showing the inter-model comparison of the percentage of pathways included on each meta pathway normalised by the total number of unique pathways per meta-pathway. The x-axis and y-axis represent the female and male data respectively. Outside a doughnut plot representing in the centre the number of total altered pathways found by gage analysis one each dataset and the percentage of pathways altered included on each meta-pathway is represented on the coronal area under each meta-pathway. The metapathways are defined in the legend.

### Proteomics analyses further support the major relevance of 16p11 gene dosage and the central role of MAPK3 and CUL3 interactors

Then, we wondered how much of the 16p11 dosage susceptible network could be confirmed by a quantitative proteomics analysis based on the hippocampus. Seven out of the 32 proteins encoded in the 16p11.2 region, were detected and successfully quantified by mass-spectrometry in the samples analysed. Their expression profiles showed a clear correlation with gene dose with lower expression in Del/+, intermediated expression in wt and Del/Dup genotype, and higher expression in Dup/+. Dup/Del values followed partially the wt abundance (Fig S5). There were 3 missing measures for Bola2, and D4A9P7 in the Del/+ group, which were very likely linked to low/borderline abundance in the samples.

Interestingly, most of the genes, contributing to the functional alteration in the 16p11 dosage and the transcription network, were also detected and quantified by the proteomic technique (highlighted in yellow with an octagonal shape (Fig. S6A). We found a strong correlation between the RNA level (RNA seq count) and our proteomic quantification in wt and Del/+ individuals for all expressed genes (Fig. S6B) and DEGs (Fig. S6C). Thus, we decided to investigate the proteomics dataset on its own and search for new insights into the 16p11 syndrome alterations. We built the proteomic-specific hippocampi 16p11 MinPPINet (Fig. S7A). Very well-connected 16p11 region proteins, such as ALDOA, BOLA2, CDIPT and COROA1, unravelled the existence of two main subnetworks (Fig. S7B): the first was around MAPK3 and SRC, a proto-oncogene coding for a membrane-bound non-receptor tyrosine kinase, while the second was built around the ATP citrate lyase (ACLY). ACLY is associated through the proteasome subunit, alpha type, 4 (PSMA4), to superoxide dismutase 1 (SOD1) and CUL3 for polyubiquitination and degradation of specific protein substrates.

### Sexual dysmorphism observed at the transcriptomics level in 16p11 LE deletion models

Then we wonder if the rat’s genetic background can also change major transcriptomics outcomes and if any sexual dysmorphism can be detected. Thus, we isolated hippocampi from 5 females and 5 males Long Evans Del/+ and controlled wild-type littermates to carry out transcriptome analysis. The DEA analysis using FCROS identified 1068 and 1324 genes specifically dysregulated (DEGs) in Del/+ males and females respectively (Table 1). Moreover, 1544 DEGs were altered independently of the sex when we ran the analysis pooling both sexes together (Fig. 8A). The computed PCA and the Euclidian distance matrix clustered the samples by their genotypes and then sex (Fig. 8B). Moreover, the specific fold change of the 16p11.2 region in both males and females showed the expected downregulation in both sexes, similar to the one observed in the male SD Del/+ male model (Fig. 8D). A good correlation occurred with the normalized counts of 16p11 region genes for wt and Del/+ genotypes in both LE male and female hippocampi (for wt R^2^=0.9996, Del/+ R^2^=0.9988) and also when comparing wt and Del/+ genotypes in males from the SD and LE genetic backgrounds (for wt R^2^=0.9966, Del/+ R^2^=0.9987) (Fig. S8). Four genes from the region were confirmed by ddPCR to follow the dosage effect with lower expression in the Del/+ rat males and females compared to wt littermates. There was a limited number of DEGs commonly deregulated in both sexes compared to hippocampal DEGs specific for males and females and more DEGS were observed in males than in females (Fig. 8C), suggesting that there was a strong influence of the sex on the genomic dysregulation induced by the deletion.

**Figure 8.**
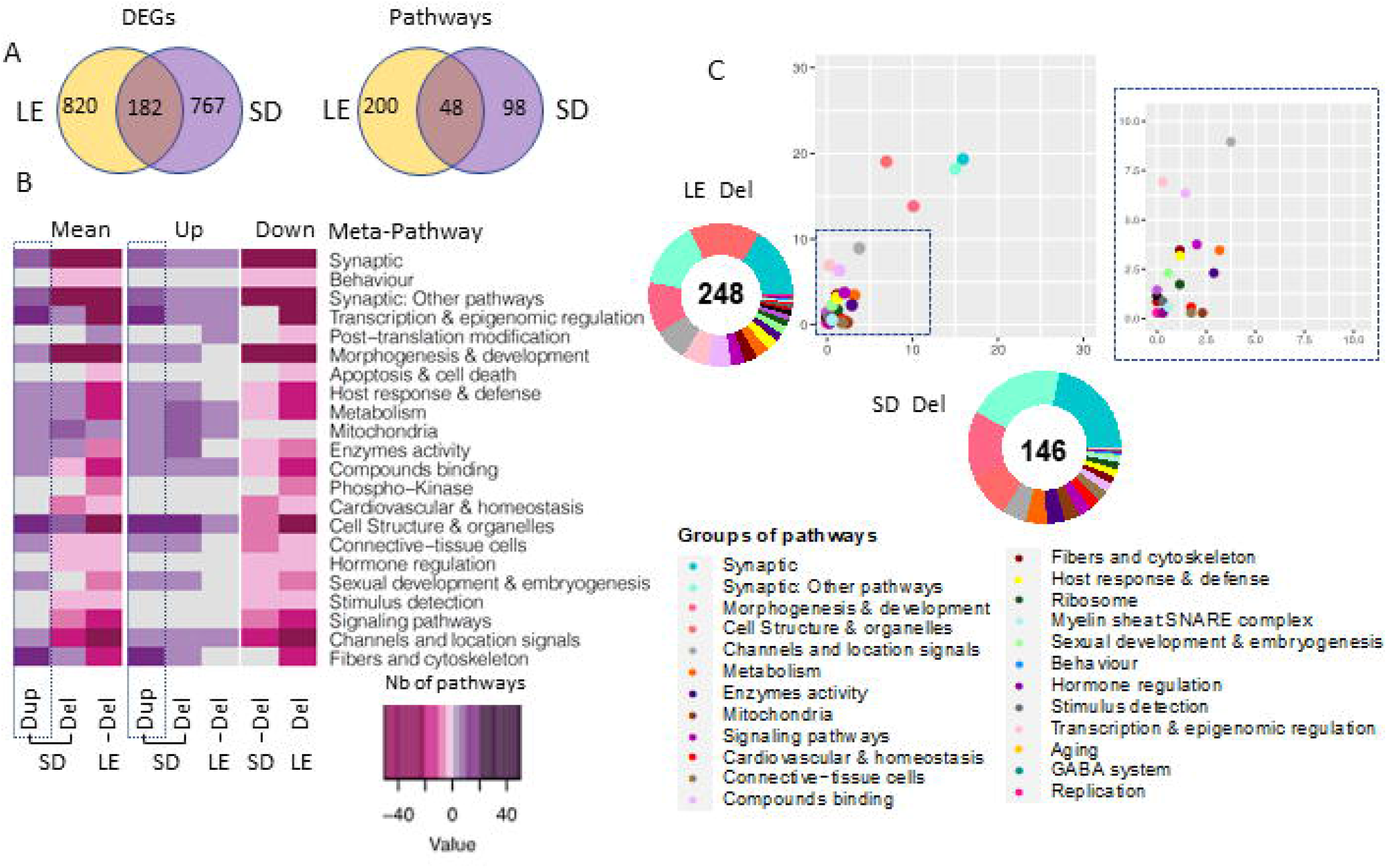
Gene expression analysis of the males LE vs SD Del (16p11) rat models. (A) Venn diagram showing in the left panel, the DEGs in common between the LE and SD male transcriptomics datasets and the background-specific DEGs. The shared DEGs correspond to a 16.2% and 20 % of the total DEGs identified respectively for the Dup and Del models. (B) Heatmap representation of the number and regulation sense of the pathways altered in the male rat from SD Del, SD Dup and LE Del models. (D) Ratio plot showing the inter-model comparison of the percentage of pathways included on each meta-pathway, normalised by the total number of unique pathways per meta-pathway. On the x-axis and y-axis represent the rat male SD Del and LE Del data respectively. Outside a doughnut plot representing in the centre the number of total altered pathways found by gage analysis for each dataset and the percentage of pathways altered included on each meta-pathway is represented on the coronal area under each meta-pathway.

Sexual dysmorphism was also found at the level of the altered pathways. With 248 pathways altered in males and 100 identified in females, only 49 specific pathways were found deregulated in both sexes but only 2 followed the same regulatory sense (Fig. 8C). We then used the classification in meta-pathways [33] to better understand the changes associated with the deletion. Looking at the resulting component, some meta-pathway like “Synaptic”, and “Synaptic: other pathways” were found downregulated in males while the resulting component was upregulated in females. Oppositely, “Behaviour”, “Host & immune response” and “Morphogenesis and development” were found downregulated in males whereas those were upregulated in females (Fig. 8E). Other meta-pathways were only found affected in Del/+ males. “Mitochondria” were only found upregulated in males while “hormone regulation” and “sexual development and embryogenesis” were downregulated only in males with no alteration in female carriers (Fig. 8E). Overall, the alteration of pathways appeared to be more pronounced in males than in females.

Looking into the number of DEGs and pathways shared and unique in both SD and LE genetic backgrounds, we identified 182 common genes, most of them following the same regulatory sense and 48 pathways, 28 upregulated and 20 downregulated in both models (Fig. 8A). Moreover, even though the number of total pathways altered in LE was higher than in SD (248 and 146 respectively), when looking at the proportion of pathways grouped on each meta-pathway considering the total number of unique pathways altered in both models only an important increase in “Cell structure” meta-pathway in LE compared to SD could be highlighted. (Fig. 8B). Considering the results obtained, we identified similar changes in the meta-pathways profiles pointing to the existence of a conserved and robust functional alteration profile (Fig. 8C), mirrored in most of the meta-pathways in the Dup/+ model. Overall the main meta-pathways for synapse (“synaptic” and “synaptic other pathways”) were commonly altered in the Del/+ models.

The few functional changes between the two genetic backgrounds were in “Apoptosis & cell death” and “post-translational modifications” which were only found affected in the LE background.

## DISCUSSION

In the present study, we described the first behavioural and cognitive phenotypes of 16p11.2 deletion and duplication of new rat models on SD and LE genetic backgrounds. A cognitive deficit was found in the novel object recognition memory test with 3 objects, and a defect in social interaction was observed with increased isolation behaviour, a typical autistic trait, in 16p11.2 *Del/+* males. The deletion of the *Sult1a1-Spn* region was also associated with the appearance of increased pinning events, a behaviour considered an expression of dominance. In addition, this type of behaviour could also be seen among pseudo-disomic *Del/Dup* carriers, suggesting a genetic construct effect not related to the dosage of genes from the region. This phenomenon may result from the new deletion allele that could alter the expression of neighbouring genes. Besides, 16p11.2 duplication in males was linked to an increase in aggressiveness. These phenotypes could be related to autistic traits and psychotic symptoms identified in patients affected by 16p11.2 rearrangements [39].

Interestingly in both outbred genetic backgrounds, the social and cognitive phenotypes were more noticeable in males than Del/+ females. The characterization of these models on a non-consanguineous genetic background allowed us to observe initially a large phenotypic variability compatible with the large symptomatic variability and the low penetrance of the neuropsychiatric disorders associated with CNVs 16p11.2 in humans. But it is important to emphasize that higher variability in the behaviour outcome of phenotypic analysis hinders our research. We have used 8 cohorts of rats to increase the number of animals (about 20-25 animals per experiment) to be able to gather a larger part of the population. For these reasons, we consider it pertinent to analyse the robustness of the phenotypes associated with the 16p11.2 deletion (CNV that has caused a more severe phenotype in the SD model) through the new rat models with outbred genetics.

While in the SD model, the variability of behaviour between individuals only allowed us to observe a trend of hyperactivity, in the new LE model we could corroborate a decrease in variability and the significant presence of hyperactivity and repetitive behaviours in males. In addition, we again detected a cognitive disorder in object recognition memory in males of the LE model, confirming the robustness of this phenotype in the 16p11.2 deletion syndrome.

Finally, when analysing the social behaviour of the LE model, we were also able to confirm the association of solitary behaviour phenotype with the deletion of the genetic interval in males. Although there were direct contact events between the tested animals, these rats avoided the behaviour of staying close to each other while exploring the test. This is a very common practice among rats, unlike mice that tend to be more solitary, which makes rats more sociable beings and animal models most useful for the study of social disorders.

In addition, curiously, we discovered that these animals spent more time approaching their test partner, which could be interpreted as cautious or scary behaviour to approach an unknown animal.

In this area also, we again observed a greater sensitivity of the male sex or a greater resilience of the female sex to the deletion of the 16p11.2 region in this new genetic background. This phenomenon is also observed in humans where more males are affected by ASD than females in the population. Our observation supports the theory of Empathy-Systematization, according to which sexual psychological differences reflect a reinforcement of systematization in the male and a reinforcement of empathy in the female. In the context of TSA, this theory has an extension, called the “extreme male brain” according to which individuals are characterized by deficiencies in empathy with an intact or increased systematization [40, 41]. Our data are also consistent with the proportion of identifying 16p11.2 rearrangements favourable for boys compared to girls reported in a previous study. This paper indicated a male: female ratio of 1.3 1 for the 16p11.2 deletion in autistic individuals and 1.6: 1 for the 16p11.2 deletion in patients with intellectual disability / developmental delay [42]. Further studies are needed for a better understanding of the mechanisms underlying risk and resilience to disease between the sexes.

Besides, we decided to evaluate the effect of 16p11.2 CNVs on the body weight of our rat models. Our study demonstrates that the deletion of the genetic interval causes only a significant reduction in body weight of young mutant males on SD background. However, for the LE model, we decided to measure the body weight of our animals at three different moments of their development. We were able to verify that the 16p11.2 deletion also causes a decrease in the weight of the mutant males at three ages, but the female sex seems to start with a normal body weight and suffer a significant loss throughout its development. These results are in line with the characterization of 16p11.2 mouse models [26]. However, in our rat model, the male rats carrying 16p11.2 duplication do not show a phenotype. On the other hand, considering the results obtained in the phenotypic study of the mouse and rat model, as opposed to the symptoms diagnosed in patients, we could hypothesize that the effect of 16p11.2 BP4-BP5 CNVs on body weight may be a specificity of the human species.

The gene expression analysis of mRNA isolated from adult rat hippocampi in SD 16p11.2 Del/+ and Dup/+ models demonstrated that 23 pathways were commonly shared and mis-regulated in both Del/+ and Dup/+. In the 16p11.2 rat models, the pathway around the primary cilium was also found altered; as described previously in mice [43]. Other changes found in the Del/+ pathways were mirrored to some extent in the Dup/+ but the severity of the changes varied between the two conditions. In addition, several additional pathways were different confirming diverse effects induced by the Del/+ and the Dup/+, as found in the mouse models [26]. One of the main alterations due to the dosage change of 16p11 was linked to the synapses, with the main central genes not linked to the regions: *Chd1, Gli1, Plg, Epha8, Disc1, Spag6l, Cfap52 or Sema3a*, except *Coro1a;* Six of which, *Chd1, Gli1, Plg, Disc1, Sema3a and Coro1a*, are reported to “abnormality of the nervous systems” in the Human Phenome Ontology. Interestingly, the most connected genes highlighted the MAPK3 and CUL3 subnetworks in the 16p11.2 models. MAPK3 is a gene from the 16p11.2 interval, thus subjected to change in dosage, and involved in the 16p11.2 syndromes [44, 45] whereas CUL3 is a target of KCTD13, another gene of the 16p11.2 region, controlling the RHOA pathway perturbed in 16p11.2 models [46, 47]. Both molecular pathways were also pointed in the proteomic studies, linked respectively with other proteins like SRC and ACLY.

Using the analysis of both sexes in the LE 16p11.2 Del model, we identified more DEGs in the male mutant hippocampi compared to female; DEGs that were also found in the SD genetic background. Of the 248 pathways altered in the Del/+ males and 100 identified in mutant females, 49 pathways were found deregulated in both sexes. More effects were observed in males than in females in various pathways, including the synapse, the Behaviour, and mitochondria. By introducing the known gene/phenotype associations, described in the DisGeNet, GO and Kegg databases, we identified *Prrt2* as a candidate gene involved in “aggressively and stereotyped behaviour” that was found upregulated in the Dup/+ model and downregulated in Del/+. We also found 4 other genes from the 16p11.2 region involved in autistic behaviour (defined by increased time in isolation) downregulated in Del and upregulated in Dup/+: *Taok2, Kctd13, Sez6l2, Mapk3*. The last two are also found in the proteomics analysis as differentially quantified peptides (Eps). Similarly, we found several genes linked to increasing time in isolation only upregulated in the Del/+ model as *Glp1r, Sema3a* or *Disc1*. Next, we wondered if we could identify any gene potentially responsible for the hypoactivity phenotype observed in the Dup model and we found 9 genes: *Coro1a*, Kctd13, Sez6l2, Spn, Aldoa, Mapk3*, Cdh1, Doc2a* and *Prrt2*. Finally, we identified 4 genes, namely *Eps, Prrt2, Mapk3*, and *Cdh1*, linked to memory and cognition deficits observed in the Del/+ model carriers and with mirroring regulatory sense in Dup/+ individuals.

Overall, the two new rat models for the 16p11.2 syndromes described here are promising in terms of behaviour alteration with more social phenotypes and similar molecular pathways, MAPK3 and KCTD13/CUL3/RHOA affected in the rat brain compared to the mouse. We already described some craniofacial changes in the SD 16p11.2 models close to the human features [48]. Nevertheless, further explorations are needed to explore the variety of phenotypes related to humans as it is currently done in the mouse. More in-depth social behaviour analysis [49] provides a more detailed description of social impairment and a strong quantitative approach is crucial to pursue if we wish one day the testing of a drug that can mitigate the social impairment observed in the 16p11.2 syndromes.

## Conflict of Interest

The authors declare that the research was conducted in the absence of any commercial or financial relationships that could be construed as a potential conflict of interest.

## Author Contribution

Conceptualization: YH; Data Curation: SML, MDM3, RW; Formal Analysis: MDM3, VN, RW; Funding Acquisition: YH, LN; Investigation: SML, AH, MP, LL, LN, VN; Methodology: JPC, LT, SM, IA, MCB, YH; Project Administration: MCB, MCP, YH; Resources: SM, LT, GP, IA, RW; Software: MDM3, RW; Supervision IA, JPC, LN, CMB, YH; Validation; SM, GP, IA, MCB, LN, RW, YH; Visualization: SML, MDM3, RW, YH; Writing – Original Draft Preparation: SML, MDM3, YH; Writing – Review & Editing: ALL.

## Funding

This work was supported by a grant from the Simons Foundation (SFARI 548888 to YH) and by the National Centre for Scientific Research (CNRS), the French National Institute of Health and Medical Research (INSERM), the University of Strasbourg (Unistra), French government funds through the “Agence Nationale de la Recherche” in the framework of the Investissements d’Avenir program by IdEx Unistra (ANR-10-IDEX-0002), a SFRI-STRAT’US project (ANR 20-SFRI-0012), ‘‘TEFOR’’ «Investissements d’Avenir» (ANRIIINSB-0014) and EUR IMCBio (ANR-17-EURE-0023) and INBS PHENOMIN (ANR-10-IDEX-0002-02) and also provided to the GenomEast platform, a member of the ‘France Génomique’ consortium for the RNASeq processing (ANR-10-INBS-0009), and to the Proteomic platform of IGBMC, that was supported by an ARC foundation grant (Orbitrap) and a Canceropole Grand Est foundation grant.. The funders had no role in the study design, data collection and analysis, decision to publish, or preparation of the manuscript.

## Acknowledgements

We would like to thank the members of the research group, of the IGBMC laboratory and of the ICS for their help in brain morphometric analysis. We extend our thanks to the animal caretakers of the ICS who are in charge of the mice well-being,

## DATA AVAILABILITY STATEMENT

The datasets for this study can be found at GEO resource database (No. GSE224935 for SD dataset and GSE for LE dataset).

## Notes

### Competing Interest Statement

The authors have declared no competing interest.

## REFERENCES

1. Redaelli S, Maitz S, Crosti F, Sala E, Villa N, Spaccini L, et al. Refining the Phenotype of Recurrent Rearrangements of Chromosome 16. International Journal of Molecular Sciences. 2019;20(5). doi: 10.3390/ijms20051095. PubMed PMID: WOS:000462542300094.

2. Hastings PJ, Lupski JR, Rosenberg SM, Ira G. Mechanisms of change in gene copy number. Nat Rev Genet. 2009;10(8):551–64. doi: 10.1038/nrg2593. PubMed PMID: 19597530; PubMed Central PMCID: PMCPMC2864001.

3. Torres F, Barbosa M, Maciel P. Recurrent copy number variations as risk factors for neurodevelopmental disorders: critical overview and analysis of clinical implications. Journal of Medical Genetics. 2016;53(2):73–90. doi: 10.1136/jmedgenet-2015-103366. PubMed PMID: WOS:000368502700001.

4. Cooper GM, Coe BP, Girirajan S, Rosenfeld JA, Vu TH, Baker C, et al. A copy number variation morbidity map of developmental delay. Nat Genet. 2011;43(9):838–46. doi: 10.1038/ng.909. PubMed PMID: 21841781; PubMed Central PMCID: PMCPMC3171215.

5. Fernandez BA, Roberts W, Chung B, Weksberg R, Meyn S, Szatmari P, et al. Phenotypic spectrum associated with de novo and inherited deletions and duplications at 16p11.2 in individuals ascertained for diagnosis of autism spectrum disorder. Journal of Medical Genetics. 2010;47(3):195–203. doi: 10.1136/jmg.2009.069369. PubMed PMID: WOS:000275771600009.

6. Marshall CR, Noor A, Vincent JB, Lionel AC, Feuk L, Skaug J, et al. Structural variation of chromosomes in autism spectrum disorder. American Journal of Human Genetics. 2008;82(2):477–88. doi: 10.1016/j.ajhg.2007.12.009. PubMed PMID: WOS:000253223900019.

7. Sanders SJ, Ercan-Sencicek AG, Hus V, Luo R, Murtha MT, Moreno-De-Luca D, et al. Multiple recurrent de novo CNVs, including duplications of the 7q11.23 Williams syndrome region, are strongly associated with autism. Neuron. 2011;70(5):863–85. doi: 10.1016/j.neuron.2011.05.002. PubMed PMID: 21658581; PubMed Central PMCID: PMCPMC3939065.

8. Steinman KJ, Spence SJ, Ramocki MB, Proud MB, Kessler SK, Marco EJ, et al. 16p11.2 Deletion and Duplication: Characterizing Neurologic Phenotypes in a Large Clinically Ascertained Cohort. American Journal of Medical Genetics Part A. 2016;170(11):2943–55. doi: 10.1002/ajmg.a.37820. PubMed PMID: WOS:000388195300022.

9. Weiss LA, Shen Y, Korn JM, Arking DE, Miller DT, Fossdal R, et al. Association between microdeletion and microduplication at 16p11.2 and autism. New England Journal of Medicine. 2008;358(7):667–75. doi: 10.1056/NEJMoa075974. PubMed PMID: WOS:000253127700003.

10. Reinthaler EM, Lal D, Lebon S, Hildebrand MS, Dahl HH, Regan BM, et al. 16p11.2 600 kb Duplications confer risk for typical and atypical Rolandic epilepsy. Hum Mol Genet. 2014;23(22):6069–80. doi: 10.1093/hmg/ddu306. PubMed PMID: 24939913.

11. Shinawi M, Liu P, Kang S-HL, Shen J, Belmont JW, Scott DA, et al. Recurrent reciprocal 16p11.2 rearrangements associated with global developmental delay, behavioural problems, dysmorphism, epilepsy, and abnormal head size. Journal of Medical Genetics. 2010;47(5):332–41. doi: 10.1136/jmg.2009.073015. PubMed PMID: WOS:000277363500005.

12. Zufferey F, Sherr EH, Beckmann ND, Hanson E, Maillard AM, Hippolyte L, et al. A 600 kb deletion syndrome at 16p11.2 leads to energy imbalance and neuropsychiatric disorders. Journal of Medical Genetics. 2012;49(10):660–8. doi: 10.1136/jmedgenet-2012-101203. PubMed PMID: WOS:000309961000009.

13. Angelakos CC, Watson AJ, O’Brien WT, Krainock KS, Nickl-Jockschat T, Abel T. Hyperactivity and male-specific sleep deficits in the 16p11.2 deletion mouse model of autism. Autism Research. 2017;10(4):572–84. doi: 10.1002/aur.1707. PubMed PMID: WOS:000400159500001.

14. Drakesmith M, Parker GD, Smith J, Linden SC, Rees E, Williams N, et al. Genetic risk for schizophrenia and developmental delay is associated with shape and microstructure of midline white-matter structures. Translational Psychiatry. 2019;9. doi: 10.1038/s41398-019-0440-7. PubMed PMID: WOS:000459835900001.

15. McCarthy SE, Makarov V, Kirov G, Addington AM, McClellan J, Yoon S, et al. Microduplications of 16p11.2 are associated with schizophrenia. Nature Genetics. 2009;41(11):1223–U85. doi: 10.1038/ng.474. PubMed PMID: WOS:000271247600015.

16. Rees E, Walters JTR, Georgieva L, Isles AR, Chambert KD, Richards AL, et al. Analysis of copy number variations at 15 schizophrenia-associated loci. British Journal of Psychiatry. 2014;204(2):108–14. doi: 10.1192/bjp.bp.113.131052. PubMed PMID: WOS:000331488900005.

17. Steinberg S, de Jong S, Mattheisen M, Costas J, Demontis D, Jamain S, et al. Common variant at 16p11.2 conferring risk of psychosis. Mol Psychiatry. 2014;19(1):108–14. Epub 20121120. doi: 10.1038/mp.2012.157. PubMed PMID: 23164818; PubMed Central PMCID: PMCPMC3872086.

18. D’Angelo D, Lebon S, Chen Q, Martin-Brevet S, Snyder LG, Hippolyte L, et al. Defining the Effect of the 16p11.2 Duplication on Cognition, Behavior, and Medical Comorbidities. Jama Psychiatry. 2016;73(1):20–30. doi: 10.1001/jamapsychiatry.2015.2123. PubMed PMID: WOS:000367820000006.

19. Jacquemont S, Reymond A, Zufferey F, Harewood L, Walters RG, Kutalik Z, et al. Mirror extreme BMI phenotypes associated with gene dosage at the chromosome 16p11.2 locus. Nature. 2011;478(7367):97–102. doi: 10.1038/nature10406. PubMed PMID: 21881559; PubMed Central PMCID: PMCPMC3637175.

20. Walters RG, Jacquemont S, Valsesia A, de Smith AJ, Martinet D, Andersson J, et al. A new highly penetrant form of obesity due to deletions on chromosome 16p11.2. Nature. 2010;463(7281):671–U104. doi: 10.1038/nature08727. PubMed PMID: WOS:000274193900038.

21. D’Angelo D, Lebon S, Chen Q, Martin-Brevet S, Snyder LG, Hippolyte L, et al. Defining the Effect of the 16p11.2 Duplication on Cognition, Behavior, and Medical Comorbidities. JAMA Psychiatry. 2016;73(1):20–30. doi: 10.1001/jamapsychiatry.2015.2123. PubMed PMID: 26629640.

22. Chawner S, Doherty JL, Anney RJL, Antshel KM, Bearden CE, Bernier R, et al. A Genetics-First Approach to Dissecting the Heterogeneity of Autism: Phenotypic Comparison of Autism Risk Copy Number Variants. American Journal of Psychiatry. 2021;178(1):77–86. doi: 10.1176/appi.ajp.2020.20010015. PubMed PMID: WOS:000604750700011.

23. Benedetti A, Molent C, Barcik W, Papaleo F. Social behavior in 16p11.2 and 22q11.2 copy number variations: Insights from mice and humans. Genes Brain Behav. 2022;21(5):e12787. Epub 20211209. doi: 10.1111/gbb.12787. PubMed PMID: 34889032.

24. Horev G, Ellegood J, Lerch JP, Son YE, Muthuswamy L, Vogel H, et al. Dosage-dependent phenotypes in models of 16p11.2 lesions found in autism. Proc Natl Acad Sci U S A. 2011;108(41):17076–81. doi: 10.1073/pnas.1114042108. PubMed PMID: 21969575; PubMed Central PMCID: PMCPMC3193230.

25. Portmann T, Yang M, Mao R, Panagiotakos G, Ellegood J, Dolen G, et al. Behavioral abnormalities and circuit defects in the basal ganglia of a mouse model of 16p11.2 deletion syndrome. Cell Rep. 2014;7(4):1077–92. doi: 10.1016/j.celrep.2014.03.036. PubMed PMID: 24794428.

26. Arbogast T, Ouagazzal A-M, Chevalier C, Kopanitsa M, Afinowi N, Migliavacca E, et al. Reciprocal Effects on Neurocognitive and Metabolic Phenotypes in Mouse Models of 16p11.2 Deletion and Duplication Syndromes. Plos Genetics. 2016;12(2). doi: 10.1371/journal.pgen.1005709. PubMed PMID: WOS:000372554100003.

27. Chenouard V, Remy S, Tesson L, Ménoret S, Ouisse LH, Cherifi Y, et al. Advances in Genome Editing and Application to the Generation of Genetically Modified Rat Models. Front Genet. 2021;12:615491. Epub 20210420. doi: 10.3389/fgene.2021.615491. PubMed PMID: 33959146; PubMed Central PMCID: PMCPMC8093876.

28. Menoret S, De Cian A, Tesson L, Remy S, Usal C, Boule JB, et al. Homology-directed repair in rodent zygotes using Cas9 and TALEN engineered proteins. Scientific Reports. 2015;5. doi: 10.1038/srep14410. PubMed PMID: WOS:000362331700001.

29. Karp NA, Meehan TF, Morgan H, Mason JC, Blake A, Kurbatova N, et al. Applying the ARRIVE Guidelines to an In Vivo Database. PLoS Biol. 2015;13(5):e1002151. doi: 10.1371/journal.pbio.1002151. PubMed PMID: 25992600; PubMed Central PMCID: PMCPMC4439173.

30. Kilkenny C, Browne WJ, Cuthill IC, Emerson M, Altman DG. Improving bioscience research reporting: the ARRIVE guidelines for reporting animal research. PLoS Biol. 2010;8(6):e1000412. doi: 10.1371/journal.pbio.1000412. PubMed PMID: 20613859; PubMed Central PMCID: PMCPMC2893951.

31. Dembélé D, Kastner P. Fold change rank ordering statistics: a new method for detecting differentially expressed genes. BMC Bioinformatics. 2014;15:14. Epub 20140115. doi: 10.1186/1471-2105-15-14. PubMed PMID: 24423217; PubMed Central PMCID: PMCPMC3899927.

32. Love MI, Huber W, Anders S. Moderated estimation of fold change and dispersion for RNA-seq data with DESeq2. Genome Biol. 2014;15(12):550. doi: 10.1186/s13059-014-0550-8. PubMed PMID: 25516281; PubMed Central PMCID: PMCPMC4302049.

33. Duchon A, Del Mar Muñiz Moreno M, Lorenzo SM, de Souza Mps, Chevalier C, Nalesso V, et al. Multi-influential genetic interactions alter behaviour and cognition through six main biological cascades in Down syndrome mouse models. Hum Mol Genet. 2021. Epub 2021/03/09. doi: 10.1093/hmg/ddab012. PubMed PMID: 33693642.

34. Ashburner M, Ball CA, Blake JA, Botstein D, Butler H, Cherry JM, et al. Gene ontology: tool for the unification of biology. The Gene Ontology Consortium. Nat Genet. 2000;25(1):25–9. doi: 10.1038/75556. PubMed PMID: 10802651; PubMed Central PMCID: PMCPMC3037419.

35. Esling P, Lejzerowicz F, Pawlowski J. Accurate multiplexing and filtering for high-throughput amplicon-sequencing. Nucleic Acids Res. 2015;43(5):2513–24. Epub 2015/02/17. doi: 10.1093/nar/gkv107. PubMed PMID: 25690897; PubMed Central PMCID: PMCPMC4357712.

36. Lindner L, Cayrou P, Jacquot S, Birling MC, Herault Y, Pavlovic G. Reliable and robust droplet digital PCR (ddPCR) and RT-ddPCR protocols for mouse studies. Methods. 2021;191:95–106. doi: 10.1016/j.ymeth.2020.07.004. PubMed PMID: WOS:000660517600011.

37. Lorbach M, Kyriakou EI, Poppe R, van Dam EA, Noldus Lpjj, Veltkamp RC. Learning to recognize rat social behavior: Novel dataset and cross-dataset application. J Neurosci Methods. 2018;300:166–72. Epub 2017/05/08. doi: 10.1016/j.jneumeth.2017.05.006. PubMed PMID: 28495372.

38. Luo W, Friedman MS, Shedden K, Hankenson KD, Woolf PJ. GAGE: generally applicable gene set enrichment for pathway analysis. BMC Bioinformatics. 2009;10:161. Epub 2009/05/27. doi: 10.1186/1471-2105-10-161. PubMed PMID: 19473525; PubMed Central PMCID: PMCPMC2696452.

39. Niarchou M, Chawner SJRA, Doherty JL, Maillard AM, Jacquemont S, Chung WK, et al. Psychiatric disorders in children with 16p11.2 deletion and duplication. Translational Psychiatry. 2019;9. doi: 10.1038/s41398-018-0339-8. PubMed PMID: WOS:000473153900003.

40. Baron-Cohen S, Lombardo MV, Auyeung B, Ashwin E, Chakrabarti B, Knickmeyer R. Why are autism spectrum conditions more prevalent in males? PLoS Biol. 2011;9(6):e1001081. Epub 2011/06/14. doi: 10.1371/journal.pbio.1001081. PubMed PMID: 21695109; PubMed Central PMCID: PMCPMC3114757.

41. Baron-Cohen S, Knickmeyer RC, Belmonte MK. Sex differences in the brain: implications for explaining autism. Science. 2005;310(5749):819–23. doi: 10.1126/science.1115455. PubMed PMID: 16272115.

42. Polyak A, Rosenfeld JA, Girirajan S. An assessment of sex bias in neurodevelopmental disorders. Genome Med. 2015;7:94. Epub 2015/08/27. doi: 10.1186/s13073-015-0216-5. PubMed PMID: 26307204; PubMed Central PMCID: PMCPMC4549901.

43. Migliavacca E, Golzio C, Maennik K, Blumenthal I, Oh EC, Harewood L, et al. A Potential Contributory Role for Ciliary Dysfunction in the 16p11.2 600 kb BP4-BP5 Pathology. American Journal of Human Genetics. 2015;96(5):784–96. doi: 10.1016/j.ajhg.2015.04.002. PubMed PMID: WOS:000354189300009.

44. Pucilowska J, Vithayathil J, Pagani M, Kelly C, Karlo JC, Robol C, et al. Pharmacological Inhibition of ERK Signaling Rescues Pathophysiology and Behavioral Phenotype Associated with 16p11.2 Chromosomal Deletion in Mice. Journal of Neuroscience. 2018;38(30):6640–52. doi: 10.1523/jneurosci.0515-17.2018. PubMed PMID: WOS:000440550000005.

45. Pucilowska J, Vithayathil J, Tavares EJ, Kelly C, Karlo JC, Landreth GE. The 16p11.2 Deletion Mouse Model of Autism Exhibits Altered Cortical Progenitor Proliferation and Brain Cytoarchitecture Linked to the ERK MAPK Pathway. Journal of Neuroscience. 2015;35(7):3190–200. doi: 10.1523/jneurosci.4864-13.2015. PubMed PMID: WOS:000349992800031.

46. Lin GN, Corominas R, Lemmens I, Yang X, Tavernier J, Hill DE, et al. Spatiotemporal 16p11.2 protein network implicates cortical late mid-fetal brain development and KCTD13-Cul3-RhoA pathway in psychiatric diseases. Neuron. 2015;85(4):742–54. doi: 10.1016/j.neuron.2015.01.010. PubMed PMID: 25695269; PubMed Central PMCID: PMCPMC4335356.

47. Martin Lorenzo S, Nalesso V, Chevalier C, Birling MC, Herault Y. Targeting the RHOA pathway improves learning and memory in adult Kctd13 and 16p11.2 deletion mouse models. Mol Autism. 2021;12(1):1. Epub 20210113. doi: 10.1186/s13229-020-00405-7. PubMed PMID: 33436060; PubMed Central PMCID: PMCPMC7805198.

48. Qiu Y, Arbogast T, Lorenzo SM, Li H, Tang SC, Richardson E, et al. Oligogenic Effects of 16p11.2 Copy-Number Variation on Craniofacial Development. Cell Reports. 2019;28(13):3320-+. doi: 10.1016/j.celrep.2019.08.071. PubMed PMID: WOS:000487587300004.

49. Rusu A, Chevalier C, View ORCID Profile de Chaumont F, View ORCID Profile Nalesso V, View ORCID Profile Brault V, Hérault Y, et al. A 16p11.2 deletion mouse model displays quantitatively and qualitatively different behaviours in sociability and social novelty over short- and long-term observation. 2022.

50. Tang H, Zhong F, Liu W, He F, Xie H. PathPPI: an integrated dataset of human pathways and protein-protein interactions. Sci China Life Sci. 2015;58(6):579–89. Epub 2015/01/15. doi: 10.1007/s11427-014-4766-3. PubMed PMID: 25591449.

51. Szklarczyk D, Morris JH, Cook H, Kuhn M, Wyder S, Simonovic M, et al. The STRING database in 2017: quality-controlled protein-protein association networks, made broadly accessible. Nucleic Acids Res. 2017;45(D1):D362–D8. Epub 2016/10/18. doi: 10.1093/nar/gkw937. PubMed PMID: 27924014; PubMed Central PMCID: PMCPMC5210637.

