## Supplementary information for "Changes in social behaviour with alterations of MAPK3 and KCTD13/CUL3 pathways in two new outbred rat models for the 16p11.2 syndromes with autism spectrum disorders"

##### Rat line engineering and genotyping

First, sgRNA was designed, produced and tested *in vitro* in rat C6 cells by T7 endonuclease I mismatch detection by the TACGENE platform (Paris, France). The sgRNA Hamont3 9revS (cr595) targeting the following sequence: gcaatctcacgtgtctgatgg and the sgRNA sgrHavalB161forw (cr 598) targeting the following sequence: gtgacatcgtgaccctgctgg were *in vitro* transcribed with the T7 high yield kit (Biolabs, New England) in the TRIP (Platform Rat Transgenesis Immunophenomic, Nantes, France) platform and were purified using EZNA microelute. These two sgRNA (20ng/µl) and Cas9 protein (3µM) were incubated at room temperature for 10 min to allow formation of RNP complexes and then were kept at 4°C until microinjection. Process and microinjection were performed as previously described (MENORET *et al.* 2015). Briefly, zygotes were collected from pre-pubescent (4–5-week-old) donor female rats (SD/Crl, Charles River, L'Arbresle, France), superovulated by injection of pregnant mare serum gonadotropin (30 IU, i.p.; Merck GmbH, Darmstadt, Germany) and, 48h later of human chorionic gonadotropin (20IU; Merck) and then mated with fertile males. One-cell-stage fertilized embryos were collected in the TRIP platform and sequentially microinjected into the male pronucleus and into the cytoplasm. Microinjected zygotes were maintained under 5% $CO_2$ at 37°C for 2h. Surviving embryos were implanted on the same day in the oviduct of pseudo pregnant females (0.5 dpc) and allowed to develop to full term.

All the animal care and procedures performed in this study were approved by the Animal Experimentation Ethics Committee of the Pays de la Loire region, France, in accordance with the guidelines from the French National Research Council for the Care and Use of Laboratory Animals (Permit Number: CEEA-PdL-2015-692).

These models were bred on the SD outbred line with the aim of helping us to understand the variability (penetrance and expressivity) of the phenotypes associated to these syndromes in patients. No impact on viability was observed because of 16p11.2 deletion and duplication in heterozygous cross of the SD rat models. The transmission of the deletion allele or the duplication were following Mendellian ratio.

Deletion of the *Sult1a1-Spn* region, referred as *Del/+*, was confirmed by PCR (Figure 1B) using Primer Del rHamont99For: (5'-GGGCTGGCAGACTTGAA-3') and Primer Del rHavalB284 Rev: (5'-GTGCCACGATCAGCAGT-3'). Duplication of the same genetic interval, referred as *Dup/+,* was identified using Primer Dup rHamont99Rev: (5'-CGCTTTGATGCCCACTAT-3') and Primer Dup rHavalB84For: (5'-AGCTGTGATCCTCTGGTT-3'). The wild-type allele was identified using Primer rAnks3-205For: (5'-CCCCAGCCTCCC ACTTGTC-3') and Primer rAnks3-205Rev: (5'-AGGATGACTGAAATTGGTGGAC-3'). The PCR reactions gave deletion, duplication and wt products of 290 bp, 500 bp and 205 bp, respectively. All rats were genotyped by PCR using the following program: 95 °C / 5 min; 35 × (72 °C / 30 s, 95 °C / 10 s, 60 °C / 10 s), 72 °C / 3 min.

For the Long Evans models, sgRNAs were designed, produced and tested in vitro using Sureguide kit (Agilent Technologies 5190-7716) by the Genetic Engineering platform (PHENOMIN-ICS, Illkirch, France). The 5' sgRNAs gR90 (MIT specificity score) targeting the following sequence TGAGATTGATGGCCGCAAGA and the sgRNA gR85 (CAATCTCACGTGTCTGATGG) and 3' sgRNAs gR80 (CATTCATTGCTCAGAGCGGC) and gR76 (CAGCCGCTCTGAGCAATGAA) were in vitro transcribed as previously described (Birling *et al.*, 2017). These 4 sgRNAs (12 ng/µl each) and Cas9 mRNA (25 ng/µl) were microinjected as previously described (Birling *et al*., 2017). Briefly, zygotes were collected from pre-pubescent (4–5-week-old) donor female rats (Long Evans, Charles River Laboratories, L'Arbresle, France), superovulated by injection of pregnant mare serum gonadotropin (30 IU, i.p.; Synchro-Part Pmsg 500, CEVA), 48h later of human chorionic gonadotropin (20IU; CHORULON® 1500, MSD) and then mated with fertile males. One-cell-stage fertilized embryos were collected in the ICS microinjection service and sequentially microinjected into the male pronucleus and into the cytoplasm. Microinjected zygotes were maintained under 5% $CO_2$ at 37°C for 2h. Surviving embryos were implanted on the same day in the oviduct of pseudo pregnant females (0.5 dpc) and allowed to develop to full term.

The second model was bred on the Long Evans outbred genetic background with no impact on the transmission or on viability of carriers. Deletion of the *Sult1a1-Spn* region, referred as *Del/+,* was confirmed by PCR using Primer Del rHamont99For : (5'-GGGCTGGCAGACTTGAA-3') and Primer Del rHavalB284 Rev: (5'-GTGCCACGATCAGCAGT-3').

The PCR reactions gave deletion product of 360 bp. All rats were genotyped by PCR using the following program: 95 °C / 5 min; 35 × (72 °C / 30 s, 95 °C / 10 s, 60 °C / 10 s), 72 °C / 3 min.

Additionally, ddPCR with a probe located in *Kcdt13* gene (*Kcdt13* is located approximately in the middle of the region of interest) was performed (as in Birling *et al*. 2017) in order to evaluation the wild type copy number in each animal and so confirm the deletion. One copy of wt was detected in the case of a Del/+ genotype.

**Behavioural analysis**

**Open Field:** This test was used to study exploration activity. Rats were tested in an automated open field (90 x 90 x 39.5 cm) made of opaque PVC with black walls and floor (Imetronic, Pessac - France). The structure was equipped with infrared sensors for accurate location and rearing behaviors of the animal. An interface provided the formatting of signals from infrared sensors and allows communication with the computer, where the software POLY OPENFIELD v5.3.2 managed experimental data. The open field arena was divided into central and peripheral regions and was homogeneously illuminated at 15 Lux. Each animal was placed in the periphery of the open field and allowed to explore freely for 30 min. The distance travelled in total arena and in each regions of the arena, as well as, the numbers of rears were recorder over the test session.

**Object location memory (OLM) task**: This test was based on the innate preference for the novelty showed by the rodents and it was carried out in the same open field arena as previously described. On the first day, rats were habituated to the arena for 15 min at 15 Lux. On the following day, animals were submitted to a first acquisition trial during 3 min in which they were individually placed in the presence of two identical objects A (syringe or flask for SD models and cup for LE model) located at 15 cm away from one of the corners, on the northeast and northwest side of the box respectively. In the case of the LE model, the test was refined by placing a reference band on the north wall of the open field. A 3-min retention trial (second trial) was conducted 5 min later, when one of the familiar objects (right or left object) was displaced randomly to a novel location (B) on the south side. The exploration time of the two objects (when the animal's snout was directed towards the object at a distance ≤ 1 cm) was recorded during both trials. Minimum total objects exploration time was set to 3 s, and rats that did not reach this criterion during the acquisition trial or retention trial were excluded from the study. We verified that no preference was seen during the exploration of the left and right object. A recognition index (RI) was defined as (($t$B / ($t$A + $t$B)) x 100). A RI of 50% corresponds to chance level and a significantly higher RI reflects good recognition memory.

**Novel object recognition (NOR) memory task:** This test allowed to evaluate the ability to recognize previously encountered objects in murine models and like the OLM task, this test is also based on the innate preference of rodents to explore novelty. We carried out NOR in the same open field arena as previously described.

Firstly, we developed the NOR test through a protocol based on the characterization of the mouse models, although we reduced the time of the trials to adapt the test to the intelligence of the rats. In the first 3 min acquisition trial, rats were presented to two identical objects A (syringe, block, bottle or flask). The animals form SD models that were evaluated in the NOR test did not belong to the same cohort as those that were analyzed in the OLM test, therefore the objects syringe and flask, were never seen prior to the NOR test by these rats. A 3-min retention trial was conducted 3 hour later. One of the two familiar objects was randomly changed for another novel object B. Test was analyzed as for the OLM task.

The surprising good performance of the mutant individuals made us question about the simplicity of this test for an intelligent animal like the rat. For this reason, we were motivated to develop a new NOR protocol.

In this case, animals from new cohorts were presented to three different objects located (A, B, C) at the northwest, northeast and southwest corner of the arena during 3-min acquisition trial. A 3-min retention trial was conducted 3 hour later. One of the three familiar objects was randomly changed for another novel object (D). The exploration time of the three objects (when the animal's snout was directed towards the object at a distance ≤ 1 cm) was recorded during both trials. Minimum exploration time was set to 3 s, and rats that did not reach this criterion during the acquisition trial or retention trial were excluded from the study. We verified that no particular objet preference was seen during the exploration. A recognition index (RI) was defined as (($t$D / ($t$A + $t$B + $t$D)) x 100). A RI of 33.3 % corresponds to chance level and a significantly higher RI reflects good recognition memory.

**Social interaction task:** This analyze focused on the evaluation of rat social behavior by manually scoring of a battery social interactions[26] among two animals of the same sex, age and genotype, housed in different cages. The test was carried in a previously described standardized open field arena during 10 min of video recording.

**Linking gene to phenotypes**

First, we downloaded the list of experimentally validated genes known to be involved in hyperactivity or locomotion behavior from the human disease database DIsGeNET (dataset: Hyperactive behavior, C0424295 with 1263 genes), in sociability (datasets: Antisocial behavior, C0233523; Antisocial Personality Disorder, C0003431; Social Phobia, C0031572; Autistic Disorder, C0004352; Aggressive behavior, C0001807), memory and cognition (datasets: Memory impairment, C0233794; Memory dysfunction, C3887551; Memory Disorder, Spatial, C0751294; Memory, Short-Term, C0025265; Age-Related Memory Disorders, C0751292; Memory performance, C1285654; Impaired cognition, C0338656 ) and stereotypies (datasets: Repetitive compulsive behavior, C1969697; Repetitive stereotyped movement, C0038271) and annotated the genes with a high confidence ortholog in rats.

We added all the genes involved in GO genesets linked to Memory and cognition: (5 GO terms: GO:0007611, GO:0007613,GO:0007614,GO:0007616, GO:0050890), Sociability (5 GO terms: GO:0035176, GO:1990911, GO:0002118, GO:0002121, GO:0002125), Locomotion (18 GO terms: GO:0007626, GO:0008344, GO:0031987, GO:0033058, GO:0035641, GO:0040011, GO:0040012, GO:0040013, GO:0040017, GO:0043056, GO:0045475, GO:0090325, GO:0090326, GO:0090327, GO:1904059, GO:1904060, GO:0036343, GO:0061744).

**LC-MS/MS Analysis**

Protein samples were analyzed using an Ultimate 3000 nano-RSLC (Thermo Scientific, San Jose California) coupled in line with a LTQ-Orbitrap ELITE mass spectrometer via a nano-electrospray ionization source (Thermo Scientific, San Jose California).

Peptide mixtures were loaded on a C18 Acclaim PepMap100 trap-column (75 µm ID x 2 cm, 3 µm, 100Å, Thermo Fisher Scientific) for 3.5 minutes at 5 µL/min with 2% ACN, 0.1% FA in H2O and then separated on a C18 Accucore nano-column (75 µm ID x 50 cm, 2.6 µm, 150Å, Thermo Fisher Scientific) with a 90 minutes linear gradient from 5% to 35% buffer B (A: 0.1% FA in H2O / B: 99% ACN, 0.1% FA in H2O), then a 20 minutes linear gradient from 35% to 80% buffer B, followed with 5 min at 99% B and 5 minutes of regeneration at 5% B. The total duration was set to 120 minutes at a flow rate of 200 nL/min. The oven temperature was kept constant at 38°C.

The mass spectrometer was operated in positive ionization mode, in data-dependent mode with survey scans from m/z 350-1500 acquired in the Orbitrap at a resolution of 120,000 at m/z 400. The 20 most intense peaks (TOP20) from survey scans were selected for further fragmentation in the Linear Ion Trap with an isolation window of 2.0 Da and were fragmented by CID with normalized collision energy of 35%. Unassigned and single charged states were rejected.

The Ion Target Value for the survey scans (in the Orbitrap) and the MS2 mode (in the Linear Ion Trap) were set to 1E6 and 5E3 respectively and the maximum injection time was set to 100 ms for both scan modes. Dynamic exclusion was used. Exclusion duration was set to 20 s, repeat count was set to 1 and exclusion mass width was ± 10 ppm.

#### Data Analysis

Proteins were identified by database searching using SequestHT (Thermo Fisher Scientific) with Proteome Discoverer™ 2.4 software (PD2.4 OPTON-30812, Thermo Fisher Scientific) (https://www.thermofisher.com/order/catalog/product/OPTON-30812) and Uniprot Rattus Norvegicus database (release 2020_02_07, 29951 entries). Precursor and fragment mass tolerances were set at 7 ppm and 0.6 Da respectively, and up to 2 missed cleavages were allowed. Oxidation (M) was set as variable modification, and Carbamidomethylation (C) as fixed modification. Peptides were filtered with a false discovery rate (FDR) at 1%, rank 1 and proteins were identified with 1 unique peptide. For the Label-Free Quantification, the protein abundancies were calculated from the average of the peptide abundancies using the TOP N (where N = 3, the 3 most intense peptides for each protein), and only the unique peptide were used for the quantification. 3210 proteins were identified along 36 MS runs (triplicate injections for 12 initial samples).

Supplemental annotation was extracted from UCSC (RGSC 6.0/rn6, Jul. 2014) and matched to the annotation from protein identification (based on UniProt mapping using Ensrnot identifyers). According to UCSC, the target region on chr 1 region 16p11.2 (defined as positions 198110001 - 198570000) contains 32 genes, 14 of these are marked as "reviewed".

Statistical analyses

All bioinformatical and statistical analyses were performed in R (R-4.0.0, https://www.r-project.org) using the packages wrMisc, wrProteo and wrGraph (available on CRAN, https://cran.r-project.org) as well as limma (Ritchie *et al.* 2015) (available on http://bioconductor.org/). Briefly, data were median-normalized and the global similarity of samples and their technical replicates examined by PCA. The annotation from UniProt was completed with genomic coordinates obtained from UCSC (https://genome.ucsc.edu/cgi-bin/hgTables).

The distribution of successful protein-quantifications was used to investigate replicate values including missing quantifications (NA from not available) and to justify subsequent imputation of NAs. Then, data were filtered to keep the rate of imputed values low (maximum 1 imputation out of 3 values for at least one pairwise-group). The contingency tables were constructed by counting all genes outside and inside the chromosome 1 target region over all samples. Contingency tables were tested using the Fisher-exact test.

Finally, all proteins were tested for direct linearity of the expected gene-dose effect. Linear regression models were constructed for each protein passing the filtering described above. The expected dose effect was set as 1.0 for *Del/+*, 2.0 for wt and *Del/Dup* as well as 3.0 for *Dup/+* samples. The resulting slope was tested by Anova for difference to 0 and confidence intervals and initial p-values corrected to Benjamini-Hochberg FDR or by the stringent Bonferroni method.

To complement the study, gene enrichment of 186 proteins with linear effects with FDR < 0.05 were tested for functional enrichment using David (https://david.ncifcrf.gov/) (HUANG *et al.* 2009) and used to construct a protein-protein-interaction network (PPI) using String (https://string-db.org)(SZKLARCZYK *et al.* 2017) and the resulting maps displayed using Cytoscape (https://cytoscape.org/)(SHANNON *et al.* 2003).

### SUPPLEMENTARY FIGURES AND TABLES

#### Supplementary Tables

| **Transcript reference** | **Exon targeted** | **Forward Primer** | **Reverse Primer** | **Probe** |
|---|---|---|---|---|
| Kctd13-201 | **Exon 2**<br>**Exon 3** | GCCTGATTGAAGACTGTCAGC | CTGTCGGGATGAGGCATAGT | **Universal Probe Library System*** UPL 7 |
| Mapk3-202 | **Exon 7**<br>**Exon 8** | CAAACAAGCGCATCACAGTAG | ATCCAGCTCCATGTCAAAGG | **ZEN™ Double-Quenched probe** HEX fluorochrom**<br>**CAGCCACTGGTTCATCTGTCGGA** |
| Prrt2-201 | **Exon 2**<br>**Exon 3** | ATCATCCTTGCCATCCTGTC | TGCTTAAGAGCTTGGCTACTC | **ZEN™ Double-Quenched probe**HEX fluorochrome**<br>**ACATTGTGGCCTTCGCTTATGCC** |
| Sez6l2-201 | **Exon 16**<br>**Exon 17** | GTCCTTGGCACTGGTGTTTA | GAAGTCTGACTCCACGGTAATG | **ZEN™ Double-Quenched probe** HEX fluorochrom**<br>**CTTTGGCTTCTCGGGCTCTCACTC** |

**Table S1: Primers for analysis of gene expression by digital droplet PCR**

| Cross | Genotype | male | female | total |
|---|---|---|---|---|
| SD Del/+ x Dup/+ | wt | 79 | 91 | 170 |
| | Del/Dup | 64 | 68 | 132 |
| | Del/+ | 69 | 69 | 138 |
| | Dup/+ | 73 | 70 | 143 |
| LE Del/+ x wt | wt | 75 | 72 | 147 |
| | Del/+ | 56 | 44** | 100** |

**Table S2. Transmission rates observed at weaning of the *Sult1a1-Spn* deletion (Del/+) or duplication (Dup/+) alleles from the SD Del/+ x Dup/+ and LE Del/+ x wt crosses**. (** chi-squared test comparing Del/+ with wt *p* <0.01).

| Cross | Genotype (nb) | Male | Female |
|---|---|---|---|
| SD Del/+ x Dup/+ | wt (n) | 525±11 | 301±6 |
| | Del/Dup (n) | 526±13 | 309±6 |
| | Del/+ (n) | 469±10** | 299±8 |
| | Dup/+ (n) | 540±13 | 305±8 |

**Table S3. Body weight of 13-weeks old adult rat from the SD 16p11.2 Del/Dup models.** *Del/+* male carriers showed decreased body weight compared to wt littermates at weeks of age. Data are mean ± SEM. Student t-test ** *p* < 0.01.

| Sex | Behavioural variable | wt | Del/Dup | Del/+ | Dup/+ |
|---|---|---|---|---|---|
| Open field | | | | | |
| Male | Horizontal activity (m) | 127 ± 5 | 132 ± 4 | 149 ± 7 | 119 ± 6 |
| | Vertical activity (count) | 139 ± 8 | 157 ± 9 | 167 ± 12 | 130 ± 7 |
| Female | Horizontal activity (m) | 174 ± 8 | 182 ± 9 | 166 ± 4 | 141 ± 6 |
| | Vertical activity (count) | 184 ± 8 | 217 ± 11 | 177 ± 13 | 156 ± 11 |
| New Object Location (5min delay) | | | | | |
| both sexes | S1 object exploration (s) | 38 ± 3 | 40 ± 3 | 48 ± 2 | 38 ± 3 |
| | S2 non-displaced object exploration (s) | 4 ± 1 | 4 ± 1 | 5 ± 1 | 6 ± 1 |
| | S2 displaced object (s) | 15 ± 1 | 8 ± 1*** | 13 ± 1 | 11 ± 1* |
| | Recognition Index (%) | 82 ± 4§§§ | 66 ± 5§§ | 73 ± 5§§§ | 70 ± 5§§§ |
| Novel Object Recognition with 3 objects and 3h of retention | | | | | |
| | S1 object A exploration (s) | 39 ± 3 | 33 ± 2 | 45 ± 3 | 38 ± 2 |
| both sexes | S2 object A exploration (s) | 12 ± 2 | 8 ± 1 | 10 ± 1 | 11 ± 1 |
| | S2 object B exploration (s) | 18 ± 3 | 13 ± 1 | 21 ± 4 | 19 ± 2 |
| | Recognition Index (%) | 64 ± 4§§§ | 63 ± 3§§§ | 62 ± 4§§ | 63 ± 4§§ |
| Male | S1 object exploration (s) | 50 ± 3 | 41 ± 4 | 44 ± 3 | 40 ± 3 |
| | S2 familiar 1 object exploration (s) | 8 ± 1 | 7 ± 1 | 13 ± 2* | 7 ± 1 |
| | S2 familiar 2 object exploration (s) | 7 ± 1 | 6 ± 1 | 16 ± 4** | 7 ± 1 |
| | S2 new object exploration (s) | 15 ± 2 | 12 ± 2 | 14 ± 2 | 13 ± 2 |
| | Recognition Index (%) | 49 ± 3§§§ | 45 ± 4§§ | 34 ± 4 | 46 ± 4§§ |
| Female | S1 object exploration (s) | 50 ± 4 | 43 ± 5 | 53 ± 3 | 42 ± 4 |
| | S2 familiar 1 object exploration (s) | 10 ± 1 | 10 ± 1 | 12 ± 1 | 10 ± 1 |
| | S2 familiar 2 object exploration (s) | 9 ± 1 | 10 ± 1 | 9 ± 1 | 10 ± 1 |
| | S2 new object exploration (s) | 18 ± 3 | 16 ± 2 | 17 ± 2 | 14 ± 2 |
| | Recognition Index (%) | 47 ± 4§§ | 44 ± 4§§ | 45 ± 3§§§ | 42 ± 4§ |
| Social Interaction | | | | | |
| Male | Solitary | 118 ± 14 | 93 ± 11 | 190 ± 22*** | 87 ± 11 |
| | Contact | 37 ± 4 | 41 ± 4 | 38 ± 7 | 60 ± 10 |
| | Head to head | 6 ± 1 | 8 ± 1 | 8 ± 2 | 7 ± 1 |
| | Head to back | 15 ± 3 | 13 ± 2 | 25 ± 5 | 13 ± 3 |
| | Approaching | 16 ± 2 | 16 ± 1 | 17 ± 2 | 17 ± 2 |
| | Moving away | 17 ± 2 | 17 ± 1 | 19 ± 2 | 16 ± 1 |
| | Following | 12 ± 3 | 15 ± 3 | 11 ± 1 | 17 ± 4 |
| | Pinning | 0 | 2 ± 1* | 3 ± 1* | 1 |
| | Agnostic | 0 | 1 | 0 | 6 ± 2** |
| Female | Solitary | 145 ± 11 | 118 ± 8 | 154 ± 12 | 144 ± 15 |
| | Contact | 38 ± 5 | 41 ± 6 | 32 ± 4 | 38 ± 5 |
| | Head to head | 7 ± 1 | 7 ± 1 | 7 ± 1 | 6 ± 1 |
| | Head to back | 35 ± 7 | 33 ± 4 | 35 ± 5 | 44 ± 8 |
| | Approaching | 21 ± 1 | 20 ± 1 | 20 ± 1 | 21 ± 1 |
| | Moving away | 25 ± 2 | 22 ± 1 | 24 ± 2 | 21 ± 1 |
| | Following | 12 ± 2 | 12 ± 2 | 17 ± 3 | 11 ± 3 |

| | | | | |
|---|---|---|---|---|
| Pinning | 0 | 0 | 0 | 1 |
| Agnostic | 0 | 0 | 0 | 2 ± 1 |

**Table S4: Behavioral variables captured for the male rat SD 16p11.2 Del and Dup models.** Data are mean ± SEM. Mann-Witney *U* test, * $p < 0.05$, ** $p < 0.01$, ***$p < 0,001$. One Sample T. Test, $^{§}p < 0,05$, $^{§§}p < 0,01$, $^{§§§}p < 0,001$.

| Sex | Age | wt | Del/+ |
|---|---|---|---|
| Male | 2 weeks | 32 ± 1 | 26 ± 1*** |
| | 4 weeks | 81 ± 2 | 61 ± 2*** |
| | 19 weeks | 534 ± 10 | 459 ± 9*** |
| Female | 2 weeks | 29 ± 1 | 26 ± 1 |
| | 4 weeks | 71 ± 3 | 60 ± 3** |
| | 19 weeks | 326 ± 9 | 298 ± 6* |

**Table S5.** Body weight of LE 16p11.2 Del rat models. *Del/+* male carriers showed decreased body weight compared to wt littermates. Data are mean ± SEM. Student t-test ** $p < 0.01$.

| Sex | Parameter | wt | Del/+ |
|---|---|---|---|
| | Open field | | |
| Male | Horizontal activity (m) | 124 ± 6 | 152 ± 9** |
| | Vertical activity (count) | 151 ± 8 | 181 ± 12* |
| Female | Horizontal activity (m) | 154 ± 4 | 180 ± 9* |
| | Vertical activity (count) | 188 ± 9 | 211 ± 15 |
| | New object location with 5 min of retention | | |
| Male | S1 object exploration (s) | 51 ± 3 | 51 ± 2 |
| | S2 non-displaced object exploration (s) | 19 ± 1 | 19 ± 2 |
| | S2 displaced object (s) | 28 ± 2 | 30 ± 2 |
| | Recognition Index (%) | 59 ± 2$^{§§§}$ | 61 ± 3$^{§§}$ |
| Female | S1 object exploration (s) | 51 ± 3 | 53 ± 3 |
| | S2 non-displaced object exploration (s) | 18 ± 2 | 18 ± 2 |
| | S2 displaced object (s) | 27 ± 2 | 28 ± 2 |
| | Recognition Index (%) | 62 ± 3$^{§§§}$ | 62 ± 3$^{§§§}$ |
| | New object recognition with 3 objects and 3 hours of retention | | |
| Male | S1 object exploration (s) | 55 ± 3 | 60 ± 3 |
| | S2 familiar 1 object exploration (s) | 11 ± 1 | 13 ± 1 |
| | S2 familiar 2 object exploration (s) | 10 ± 1 | 12 ± 1 |
| | S2 new object exploration (s) | 17 ± 2 | 16 ± 1 |
| | Recognition Index (%) | 44 ± 3$^{§§§}$ | 39 ± 3 |
| Female | S1 object exploration (s) | 62 ± 2 | 58 ± 2 |
| | S2 familiar 1 object exploration (s) | 14 ± 1 | 12 ± 2 |
| | S2 familiar 2 object exploration (s) | 12 ± 1 | 15 ± 2 |

| | | | |
|---|---|---|---|
| | S2 new object exploration (s) | 20 ± 1 | 18 ± 1 |
| | Recognition Index (%) | 44 ± 3$^{§§§}$ | 41 ± 2$^{§§}$ |
| | Social interaction | | |
| Male | Solitary | 106 ± 14 | 140 ± 6* |
| | Contact | 64 ± 5 | 82 ± 7 |
| | Head to head | 5 ± 1 | 4 ± 1 |
| | Head to back | 10 ± 1 | 9 ± 2 |
| | Approaching | 12 ± 1 | 15 ± 1* |
| | Moving away | 15 ± 1 | 17 ± 1 |
| | Following | 10 ± 1 | 13 ± 2 |
| | Pinning | 0 | 0 |
| | Agnostic | 0 | 0 |
| Female | Solitary | 117 ± 10 | 147 ± 15 |
| | Contact | 52 ± 4 | 54 ± 4 |
| | Head to head | 4 | 4 |
| | Head to back | 29 ± 5 | 19 ± 4 |
| | Approaching | 10 ± 1 | 11 ± 1 |
| | Moving away | 12 ± | 13 ± 1 |
| | Following | 14 ± 2 | 11 ± 2 |
| | Pinning | 0 | 0 |
| | Agnostic | 0 | 0 |

**Table S6: Behavioral variables captured for the rat LE 16p11.2 Del model in both sexes.** Data are mean ± SEM. Mann-Witney *U* test, * $p < 0.05$, ** $p < 0.01$, ***$p < 0,001$. One Sample T. Test, $^{§}p < 0,05$, $^{§§}p < 0,01$, $^{§§§}p < 0,001$.

**Table S7: Correlation of the DEGs hippocampal expression level with the Copy number of the homologours region to human 16p11.2 in rat SD and LE models.** (Separated Excell document)

**Table S8 DEGs and pathways misregulared in 16p11.2 hippocampi rat models.** (Separated Excell document)

#### SUPPLEMENTARY FIGURES

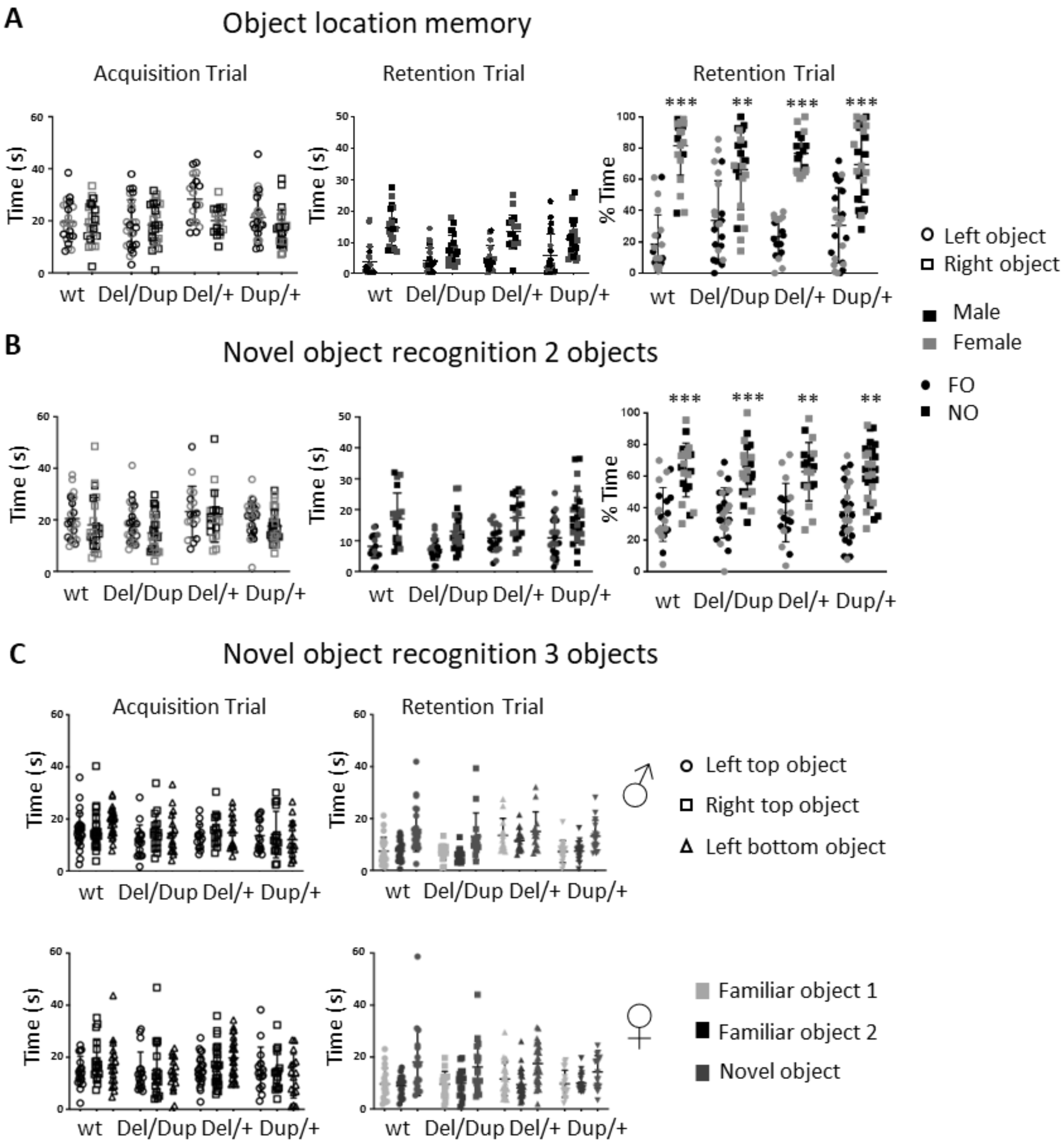


**Supplementary Figure 1: Novel object location (A), and object recognition with 2 (B) or 3 objects (C) in the 16p11.2 SD models.** (A, B) Novel object location and novel object recognition memory task with 2 objects of the 16p11.2 rat models. The left graphs shows the absolute exploration time in seconds of the left object and the right object during the acquisition trial (3 minutes) from all genotypes. The central graphs shows the absolute exploration time in seconds of the familiar position object in OLM or familiar object in NOR observed previously (FO) and the new position object in NOL or novel object in NOR (NO) during the retention trial (3 minutes) from wt and mutant rats. The right graphs shows the percentage exploration time of each object during the retention trial. In the OLM test, male (black) and female (grey) rats from different genotypes (wt (n=21), *Del/Dup* (n=25), *Del/+* (n=18) and *Dup/+* (n=25) showed increased time in exploring the NO compared to the time spent exploring the FO. In the NOR test, our rat models from all genotypes (wt (n=22), *Del/Dup* (n=25), *Del/+* (n=18) and *Dup/+* (n=27) show an exploration preference for the new object. (C) Novel object recognition memory task with 3 objects of the 16p11.2 rat models for both sex. The left graph show the absolute exploration time in seconds of the left top object, the right top object and the left bottom object during the acquisition trial (3 minutes) from all genotypes. The right graph show the absolute exploration time in seconds of the 2 familiar objects observed previously and the novel object during the retention trial (3 minutes) from our rat models. Student t-test with ** $p < 0.01$ and *** $p < 0.001$

```
                                                        Del 483 Kb (483040 pb)
                                                  Chr1: 198,100,544/198,583,584
Del  AAGGAGTTCCTATGATCCAATAAATACTCTTCCCTCTCACCATGTCTCCTGGGCCTCACTACGTCAGACCCTCTTGCCC
     TCCTCCAGACACGTGAGATTGATGGCCgctcagagcggctggccattacctccataaaggtcattctcttggcaagctg
     catttgtgtaaggcactcttagggttctctaTGCTGTCCTAGTTTGGAGAAGAACACTACATGCTTCTGGAAAATTCAC
     ATTCAAGCAGCAACATCTTAACATATCTCCTCACAGATACATTTTATATTATGTGTGAACTGCTGATCGTGGCAC
```

**Supplementary Figure 2: Sequence of the junction of the deletion in the 16p11.2 LE deletion models**. The positions are indicated with reference to the RGSC 6.0/rn6 rat genome sequence.

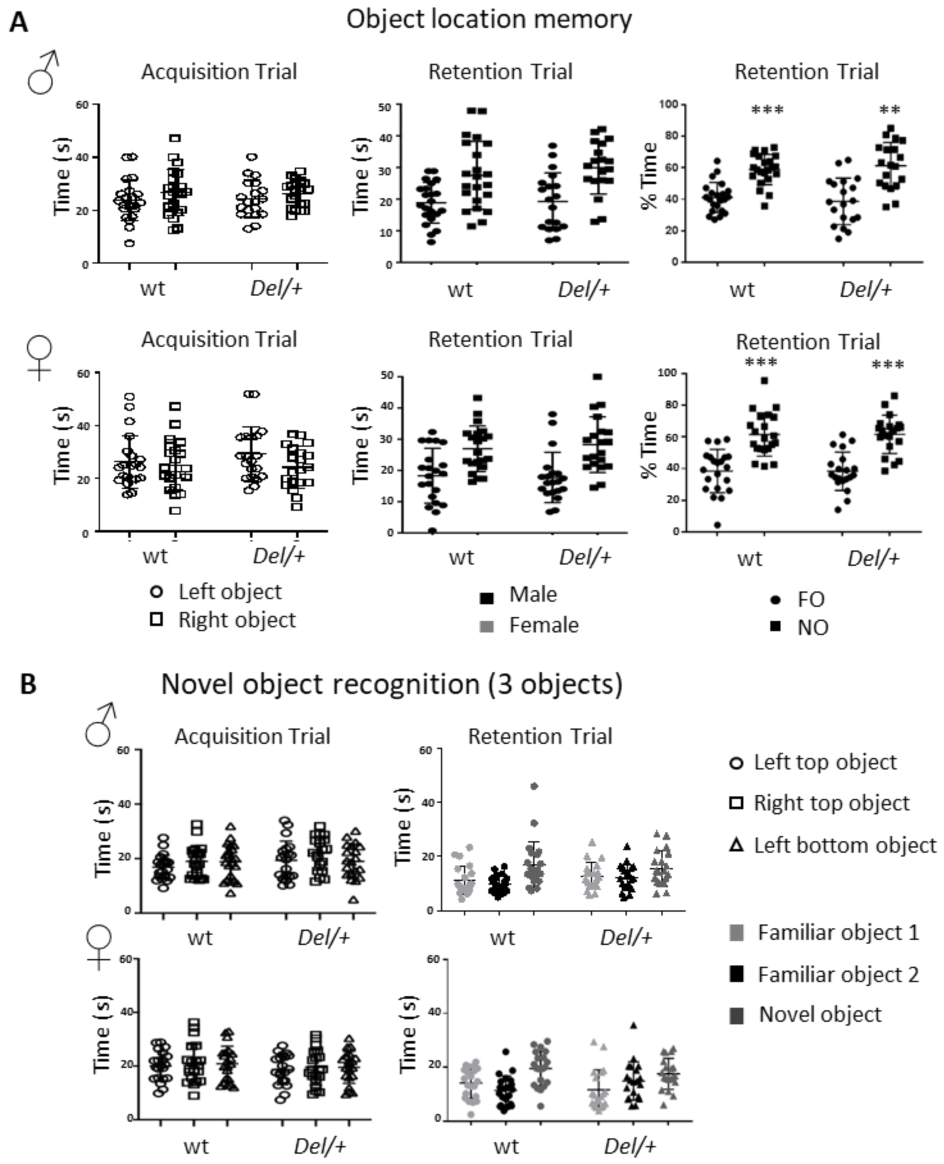


**Supplementary Figure 3: Time spent during the novel object location (A) and object recognition with 3 objects, (B) assessment in the 16p11.2 LE models.** (A) Novel object location recognition memory task with 2 objects of the LE *Del/+* 16p11.2 rat model. The left graphs shows the absolute exploration time in seconds of the left object and the right object during the acquisition trial (3 minutes). The central graphs shows the absolute exploration time in seconds of the familiar position object (FO) and the new position object (NO) during the retention trial (3 minutes) from wt and mutant rats. The right graphs shows the percentage exploration time of each object during the retention trial. Male (wt (n= 22) and *Del/+* (n= 19)) and female (wt (n= 21) and *Del/+* (n= 20)) rats from LE 16p11.2 rat model did not have their object location memory affected. The mutants of both sexes show a preference for the object whose location is new and therefore the recognition index of new location is statistically higher than 33.3% (One sample t test for males : wt ($t_{(21)} = 4.35$; $p = 0.0003$) and *Del/+* ($t_{(18)} = 3.35$; $p = 0.0036$; One sample t test for females : wt ($t_{(20)} = 3.88$; $p = 0.0009$) and *Del/+* ($t_{(19)} = 4.29$; $p = 0.0004$). (B) Novel object recognition memory task with 3 objects of the LE *Del/+* 16p11.2 rat model for both sex. The left graph show the absolute exploration time in seconds of the left top object, the right top object and the left bottom object during the acquisition trial (3 minutes) from all genotypes. The right graph show the absolute exploration time in seconds of the 2 familiar objects observed previously and the novel object during the retention trial (3 minutes) from our rat models. Student t-test with ** $p < 0.01$ and *** $p < 0.001$

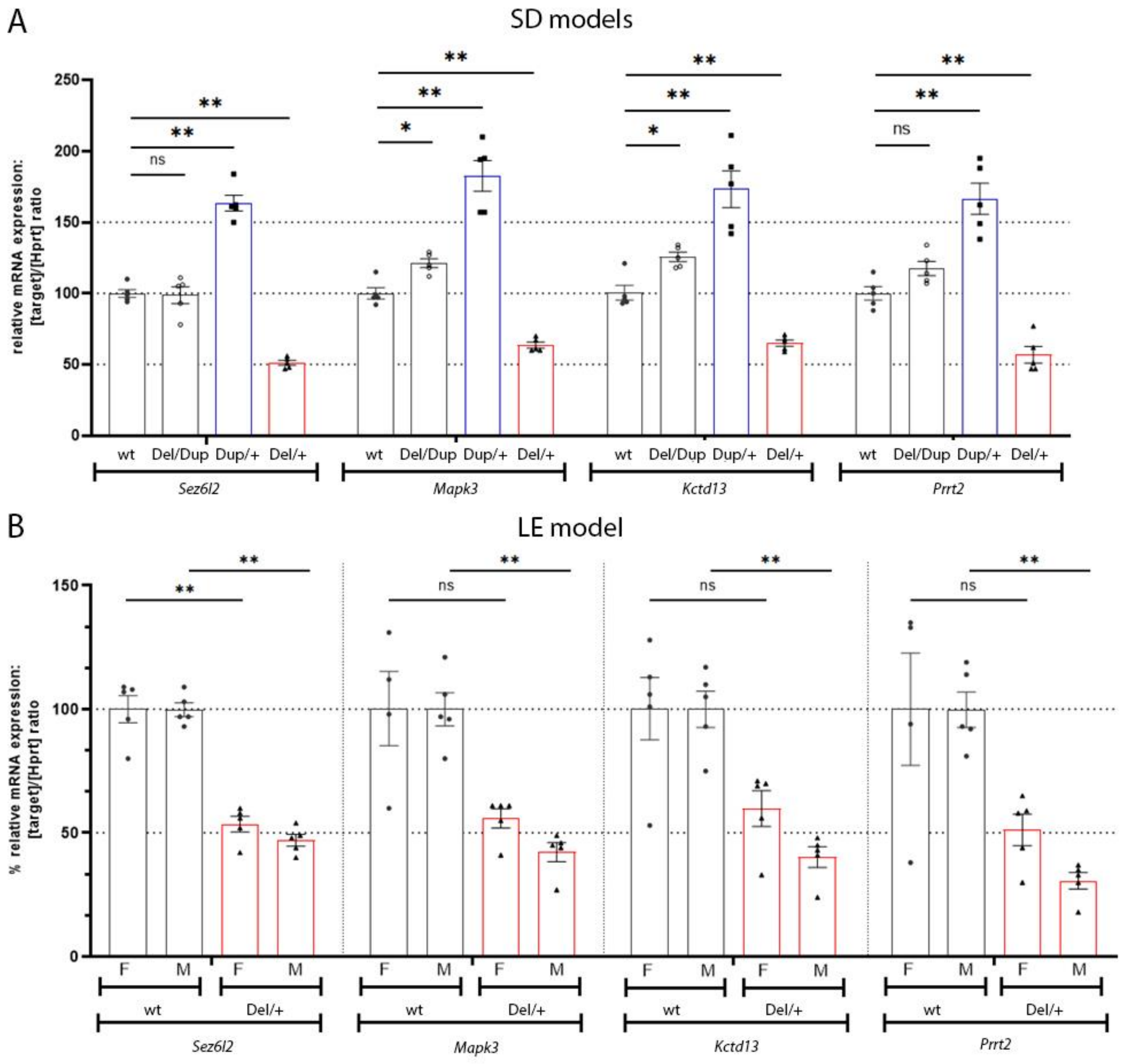

**Supplementary Figure 4: Relative mRNA expression level of genes from the 16p11.2 regions compared to Hprt control gene measured by digital droplet PCR. The** analysis was done on 5 samples derived from the hippocampi of individuals isolated from the different SD (A) and LE (B) models.

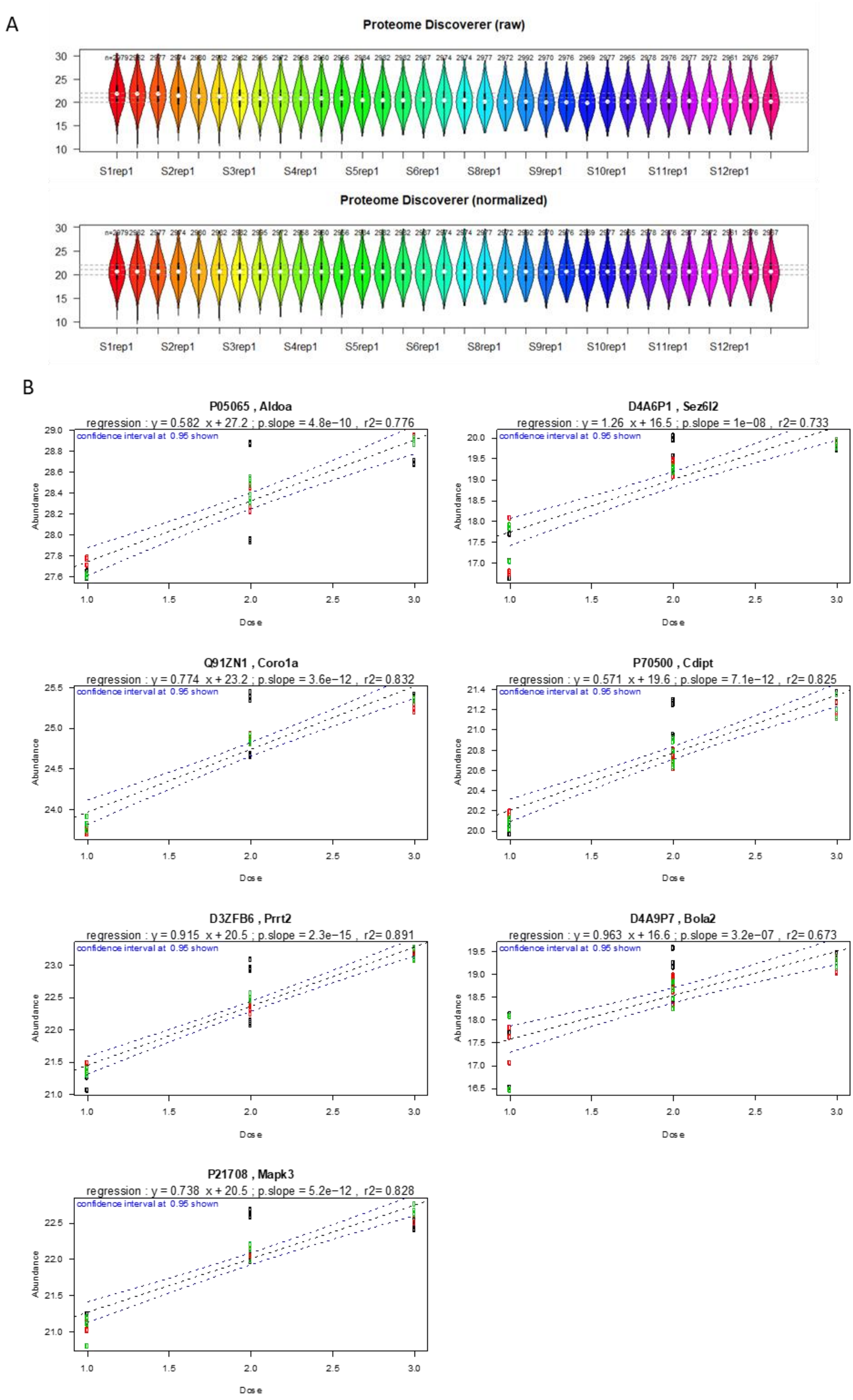
A
Proteome Discoverer (raw)
S1rep1 S2rep1 S3rep1 S4rep1 S5rep1 S6rep1 S8rep1 S9rep1 S10rep1 S11rep1 S12rep1
Proteome Discoverer (normalized)
S1rep1 S2rep1 S3rep1 S4rep1 S5rep1 S6rep1 S8rep1 S9rep1 S10rep1 S11rep1 S12rep1
B
P05065 , Aldoa
regression : y = 0.582 x + 27.2 ; p.slope = 4.8e−10 , r2= 0.776
D4A6P1 , Sez6l2
regression : y = 1.26 x + 16.5 ; p.slope = 1e−08 , r2= 0.733
Q91ZN1 , Coro1a
regression : y = 0.774 x + 23.2 ; p.slope = 3.6e−12 , r2= 0.832
P70500 , Cdipt
regression : y = 0.571 x + 19.6 ; p.slope = 7.1e−12 , r2= 0.825
D3ZFB6 , Prrt2
regression : y = 0.915 x + 20.5 ; p.slope = 2.3e−15 , r2= 0.891
D4A9P7 , Bola2
regression : y = 0.963 x + 16.6 ; p.slope = 3.2e−07 , r2= 0.673
P21708 , Mapk3
regression : y = 0.738 x + 20.5 ; p.slope = 5.2e−12 , r2= 0.828
confidence interval at 0.95 shown
Abundance
Dose

**Supplementary Figure 5: Linear correlation with genetic dosage of the abundance of 7 proteins from the 16p11 rat syntenic region detected in the proteomic analysis done from the Del/+, Dup/+, wt and Del/Dup genotypes in the 16p11.2 SD models**. The plot was drawn considering on y-axis the protein abundance and in the X-axis the samples were represented considering their 16p11 gene dosage as follows: At dose =1 Del/+ samples were represented, dose=2 the Wt and Del/Dup(16p11) samples and at dose=3 the Dup/+ samples. The regression coefficients are shown in the upper part of each plot. Inside each plot, the dotted lines denoting the confidence interval are drawn in blue. This view shows again that the data follows a linear distribution, however some deviations to linearity are present together with certain amount of spread/noise.

A

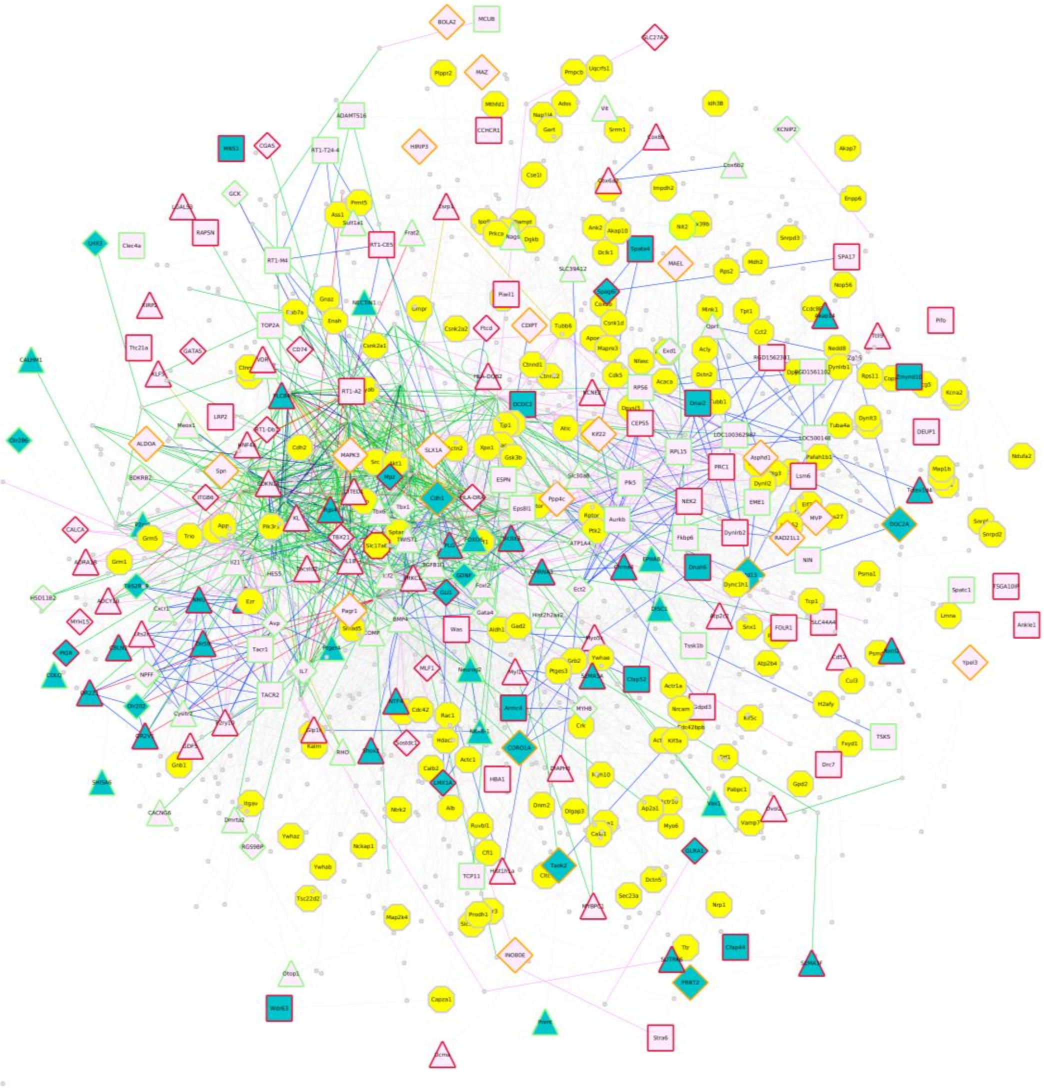

B Expressed genes found in Proteomics

Wt

Del

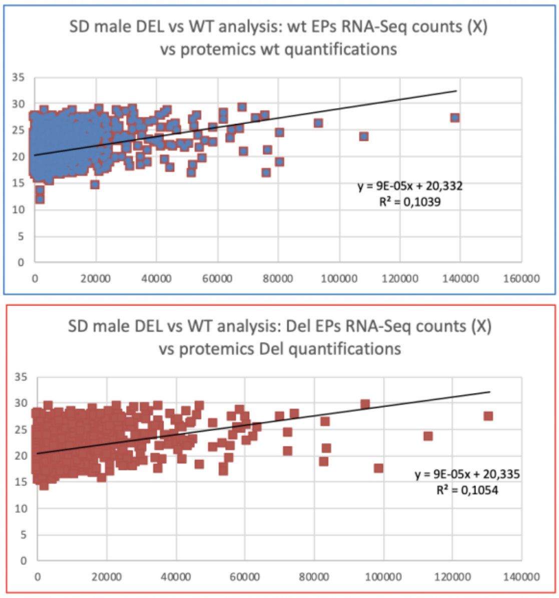


C

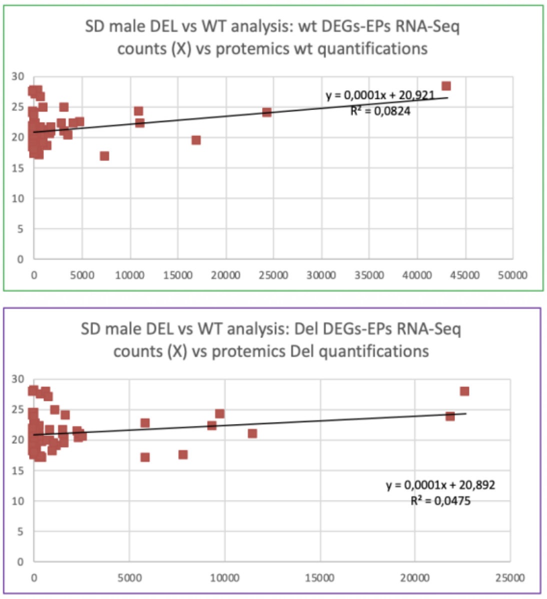

**Supplementary Figure 6: Minimun rat RegPPINet linked to synaptic function gene and expression correlation analyses in 16p11.2 SD models.** (A) genes linked to synaptic function (GO/KEGG association databases) colouring the inner shape of the nodes in green. The sense of regulation of those genes in the rat Del/+ and Dup/+ transcriptomic datasets is represented with the border color of each node as follows: if the gene was found downregulated in both models, the border is green; instead if it was found upregulated in both, the border is pink, or if the genes are found with opposing regulatory sense on the datasets, the border is orange. Highlighted in yellow within the inner node colour are the proteins successfully quantified by proteomics carried on in the Del/+ and Dup/+ rat models. The edges colour represent the type of interaction annotated by following the PathPPI classification (TANG *et al.* 2015), and ReactomeFIViz annotations as follows i) The GErel edges indicating expression were coloured in blue and repression in yellow. ii) PPrel edges indicating activation were coloured in green, inhibition in red. Iii) Interactions between proteins known to be part of complexes in violet. Iv) Predicted interactions were represented in grey including the PPI interactions identified by STRING DB (SZKLARCZYK *et al.* 2017) after merging both networks. (B) Upper panel, correlation of the mean of wt samples gene counts for all the rat expressed genes (notes as EGs) identified in the transcriptomics represented on the x-axis and the protein abundance quantified in the rat SD Del/+ proteomics datasets. Bottom panel, correlation of the mean of Del/+ samples gene counts for all the rat expressed genes (notes as EGs) identified on the transcriptomics represented in the x-axis and the protein abundance quantified in the rat Del/+ proteomics datasets. **(**C) Upper panel, correlation of the mean of wt samples gene counts for all the rat differentially expressed genes (notes as DEGs) identified on the transcriptomics represented on the x-axis and the protein abundance quantified in the mouse Del/+ proteomics datasets. Bottom panel, correlation of the mean of Del gene counts for the rat differentially expressed genes (notes as DEGs) identified on the transcriptomics represented on the x-axis and the protein abundance quantified in the rat Del/+ proteomic datasets.

A

B

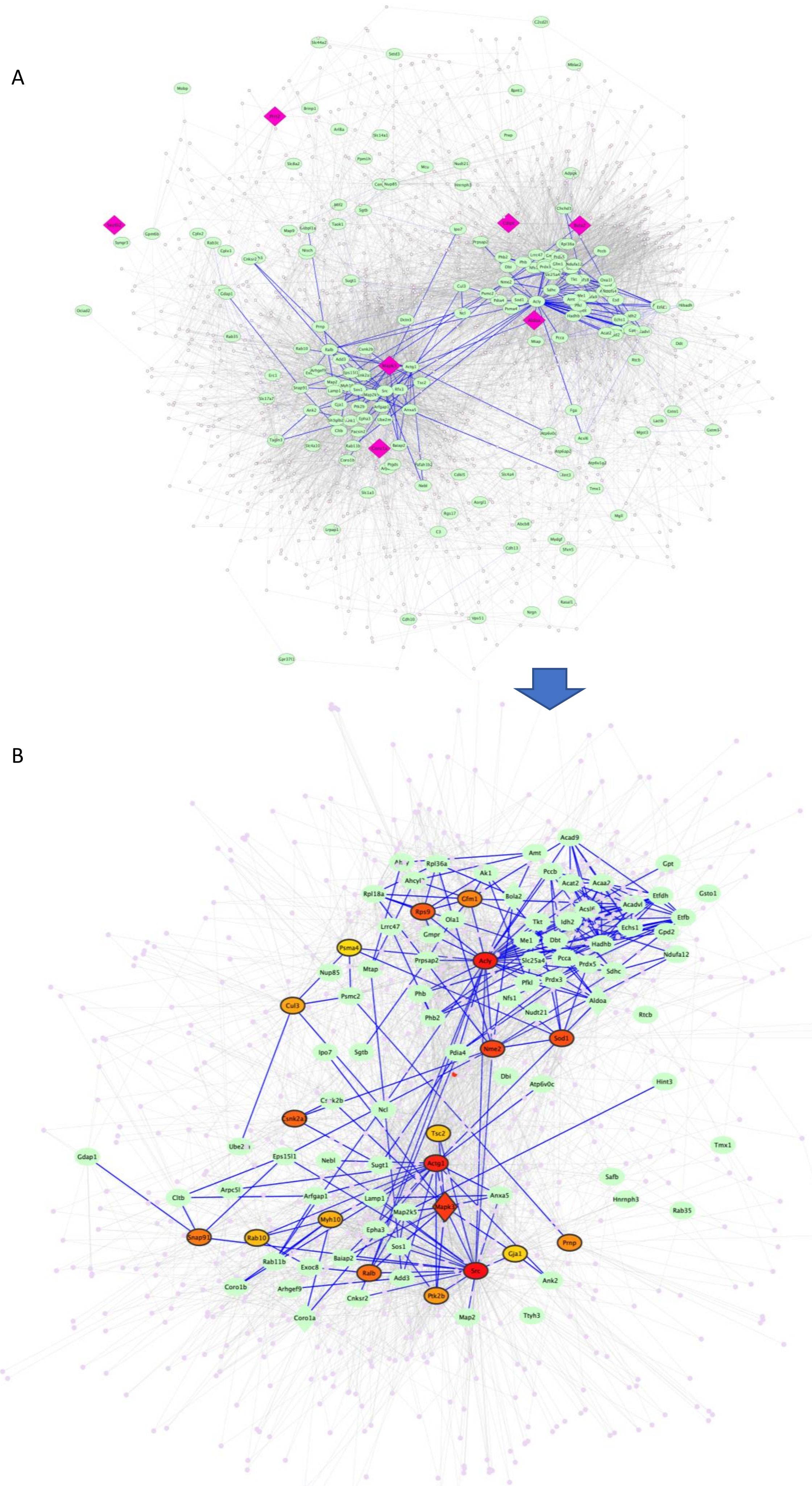

Graph legend
Central hubs
by Betweenes

| Rank | Protein |
|---|---|
| 1 | Src |
| 2 | Acly |
| 3 | Actg1 |
| 4 | Ubb |
| 5 | Mapk3 |
| 6 | Nme2 |
| 7 | Sod1 |
| 8 | Rps9 |
| 9 | Csnk2a1 |
| 10 | Ralb |
| 11 | Snap91 |
| 12 | Gfm1 |
| 13 | Prnp |
| 14 | Ptk2b |
| 15 | Cul3 |
| 16 | Myh10 |
| 17 | Rab10 |
| 18 | Tsc2 |
| 19 | Gja1 |
| 20 | Psma4 |
| | Not central seeds |
| | Connecting seeds |

**Supplementary Figure 7: proteomic based PPI network derived from SD Del and Dup male hippocampi.** (A) Twenty central nodes of the altered proteomics network in the Del and Dup 16p11 SD male hippocampi identified by Betweenes index. The 197 altered DQPs (differentially quantified peptides) seeds were used to build a Minimun PPI Network using String Db and Network analysis: 177 of those were annotated with at least one PPI using STRING with a medium confidence level of 400 and accepting PPIs identified using all sources of annotation. The network visualization was organised based on the Edge-weighted spring embedded by betweenness index layout. Graph legend, the edges connecting the DQPs seeds are coloured in blue, instead other PPIs in grey**.** (B) Highlighting in yellow the nodes found also expressed in the transcriptomics SD and LE datasets. The top 20 central network have 743 nodes and 4011 edges of those, 106 are seeds, 7 are 16p11 region genes. Moreover 600 QPs were also identified in the transcriptomics analyses with a mean expression higher than 100, (569 higher than 200, 544 higher than 300, 486 higher than 600, 449 higher than 800) of those 61 were identified as DEGs in the transcriptomics datasets for LE or SD.

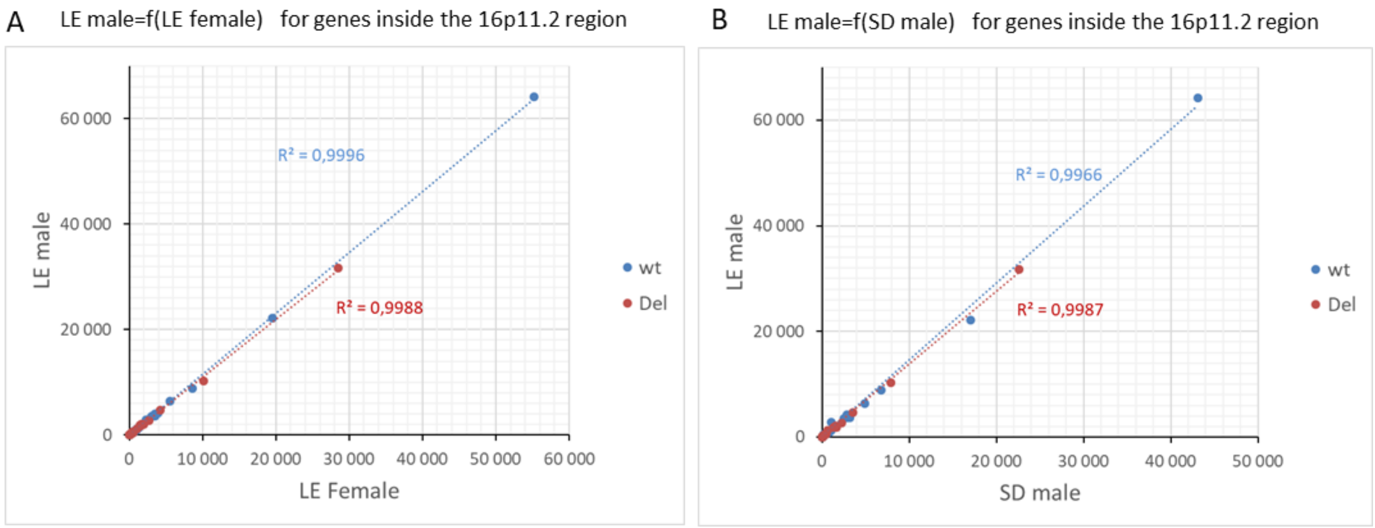


**Supplementary Figure 8: Correlation of hippocampal gene expression for the genes of the 16p11 interval with the genes in wt and Del genotype in the rat 16p11.2 models.** (A) correlation of the mean of gene expression between female and male from the LE genetic background. (B) correlation of the mean of gene expression between males from the SD and LE genetic backgrounds represented on the y-axis.

**REFERENCES to Supplementary information**


Huang, d. W., B. T. Sherman and R. A. Lempicki, 2009 Systematic and integrative analysis of large gene lists using DAVID bioinformatics resources. Nat Protoc 4**:** 44-57.

Menoret, S., A. De Cian, L. Tesson, S. Remy, C. Usal *et al.*, 2015 Homology-directed repair in rodent zygotes using Cas9 and TALEN engineered proteins. Scientific Reports 5.

Ritchie, M. E., B. Phipson, D. Wu, Y. Hu, C. W. Law *et al.*, 2015 limma powers differential expression analyses for RNA-sequencing and microarray studies. Nucleic Acids Res 43**:** e47.

Shannon, P., A. Markiel, O. Ozier, N. S. Baliga, J. T. Wang *et al.*, 2003 Cytoscape: a software environment for integrated models of biomolecular interaction networks. Genome Res 13**:** 2498-2504.

Szklarczyk, D., J. H. Morris, H. Cook, M. Kuhn, S. Wyder *et al.*, 2017 The STRING database in 2017: quality-controlled protein-protein association networks, made broadly accessible. Nucleic Acids Res 45**:** D362-D368.

Tang, H., F. Zhong, W. Liu, F. He and H. Xie, 2015 PathPPI: an integrated dataset of human pathways and protein-protein interactions. Sci China Life Sci 58**:** 579-589.
